# Immunization with a self-assembling nanoparticle vaccine displaying EBV gH/gL protects humanized mice against lethal viral challenge

**DOI:** 10.1101/2022.02.17.480914

**Authors:** Harman Malhi, Leah J. Homad, Yu-Hsin Wan, Bibhav Poudel, Brooke Fiala, Andrew J. Borst, Jing Yang Wang, Carl Walkey, Jason Price, Abigail Wall, Suruchi Singh, Zoe Moodie, Simran Handa, Colin Correnti, Barry L. Stoddard, David Veesler, Marie Pancera, James Olson, Neil P. King, Andrew T. McGuire

**Affiliations:** Vaccine and Infectious Disease Division, Fred Hutchinson Cancer Research Center, Seattle WA; Department of Biochemistry, University of Washington, Seattle, WA; Institute for Protein Design, University of Washington, Seattle, WA; Clinical Research Division, Fred Hutchinson Cancer Research Center Seattle, WA; Division of Basic Sciences, Fred Hutchinson Cancer Research Center, Seattle, WA; Howard Hughes Medical Institute; Department of Global Health, University of Washington, Seattle, WA; Department of Laboratory Medicine and Pathology, University of Washington, Seattle WA

## Abstract

Epstein-Barr virus (EBV) is a cancer-associated pathogen responsible for 140,000 deaths per year. EBV is also the etiological agent of infectious mononucleosis and is associated with multiple sclerosis and rheumatoid arthritis. Thus, an EBV vaccine could alleviate significant morbidity and mortality. EBV is orally transmitted and has tropism for both epithelial cells and B cells which are present in the oral cavity. Therefore, a prophylactic vaccine would need to prevent infection of both cell types. Passive transfer neutralizing monoclonal antibodies targeting the viral gH/gL glycoprotein complex prevent experimental EBV infection in humanized mice and rhesus macaques, suggesting that gH/gL is an attractive vaccine candidate. Here, we produced and evaluated the immunogenicity of several nanoparticle immunogens displaying gH/gL with distinct valencies and geometries. After one or two immunizations, all nanoparticles elicited superior binding and neutralizing titers relative to monomeric gH/gL. Antibodies elicited by a computationally designed self-assembling nanoparticle that displays 60 copies of the gH/gL protein conferred protection against a lethal dose of EBV in a humanized mouse challenge model, whereas antibodies elicited by monomeric gH/gL did not. Taken together, these data motivate further development of gH/gL nanoparticle vaccine candidates for EBV.

## Introduction

Epstein-Barr virus (EBV) is one of the most common human viruses. It is a herpesvirus with tropism for both B cells and epithelial cells and is associated with several malignancies of these two cell types including Hodgkin lymphoma, Burkitt lymphoma, diffuse large B cell lymphoma, post-transplant lymphoproliferative disease, nasopharyngeal carcinoma and gastric carcinoma (Cohen, 2018; Kutok and Wang, 2006; Shannon-Lowe and Rickinson, 2019; Taylor et al., 2015). It is estimated that EBV is responsible for ∼200,000 new cases of cancer and ∼140,000 cancer deaths globally per year (Cohen et al., 2011; Khan and Hashim, 2014; Shannon-Lowe and Rickinson, 2019). EBV is also the causative agent of infectious mononucleosis (IM) and is linked to multiple sclerosis and rheumatoid arthritis (Angelini et al., 2013; Balandraud and Roudier, 2018; Bjornevik et al., 2022; Handel et al., 2010; Levin et al., 2010; Munger et al., 2011; Thacker et al., 2006). Thus, a vaccine that prevents EBV infection and/or associated pathologies would have a significant global health impact (Ainsworth, 2018; Cohen *et al*., 2011; Shannon-Lowe and Rickinson, 2019).

EBV is orally transmitted and both B cells and epithelial cells are present in the oropharynx. Thus, an effective vaccine would likely need to prevent or severely limit infection in both cell types (Rickinson et al., 2014; Taylor *et al*., 2015). The dual tropism of EBV infection is accomplished through the orchestrated function of multiple glycoproteins (Connolly et al., 2021). gH, gL and gB constitute the core fusion machinery and are essential for viral entry irrespective of cell type. gB is a transmembrane fusion protein that promotes the merger of the viral and host membranes (Backovic et al., 2009). gB activity depends on the heterodimeric gH/gL complex, which is essential for infection and regulates fusion (Haddad and Hutt-Fletcher, 1989; Mohl et al., 2016; Oda et al., 2000; Stampfer and Heldwein, 2013). Epithelial cell infection is initiated by the binding of the viral BMRF-2 protein to β1 integrins on the cell surface (Tugizov et al., 2003). Following attachment, binding of gH/gL to one or more cell surface receptors is thought to induce a conformational change that triggers gB activation. αvβ6, and αvβ8 integrins, neuropilin 1, non-muscle myosin heavy chain IIA and the ephrin A2 receptor have all been implicated as gH/gL receptors (Chen et al., 2018; Chesnokova et al., 2009; Su et al., 2020; Wang et al., 2015; Xiong et al., 2015; Zhang et al., 2018).

Viral attachment to B cells is mediated by gp350, which binds to complement receptors (CR) 1 and 2 (Nemerow et al., 1987; Ogembo et al., 2013; Tanner et al., 1987). The triggering of gB during B cell entry depends on the tripartite complex of gH/gL and the vial glycoprotein gp42. Binding of gp42 to the B chain of human leukocyte antigen class II leads to activation of gB through the gH/gL/gp42 complex (Haan et al., 2000; Sathiyamoorthy et al., 2014; Spriggs et al., 1996).

Neutralizing antibodies are the correlate of protection for most effective vaccines (Gilbert et al., 2022; Plotkin, 2010). It is therefore likely that they will be an important component of an immune response elicited by an EBV vaccine. Serum from naturally infected individuals can neutralize EBV infection of B cells and epithelial cells (Miller et al., 1972; Moss and Pope, 1972; Sashihara et al., 2009; Tugizov *et al*., 2003), and all the viral proteins involved in viral entry are targeted by neutralizing antibodies (Bu et al., 2019; Thorley-Lawson and Poodry, 1982; Tugizov *et al*., 2003; Xiao et al., 2009). To date, most EBV subunit vaccine efforts have focused on gp350. gp350 is capable of adsorbing most of the serum antibodies that neutralize EBV infection of B cells (Bu *et al*., 2019; Thorley-Lawson and Poodry, 1982).

Mechanistically, neutralizing anti-gp350 monoclonal antibodies (mAbs) block the gp350-CR1/CR2 interaction (Hoffman et al., 1980; Mutsvunguma et al., 2019; Ogembo *et al*., 2013; Szakonyi et al., 2006; Tanner et al., 1988). However, antibodies against gp350 are ineffective at inhibiting EBV infection of CR^-^ epithelial cells and can enhance infection of this cell type (Molesworth et al., 2000; Tugizov *et al*., 2003; Turk et al., 2006). Passive transfer of a neutralizing anti-gp350 mAb protected one of three macaques against high-dose experimental infection with rhesus lymphocryptovirus, the EBV ortholog that infects macaques (Mühe et al., 2021) indicating that gp350 antibodies could be protective *in vivo.* A phase II trial of a gp350 vaccine failed to protect against EBV despite decreasing the incidence of symptomatic infectious mononucleosis by 78% (Sokal et al., 2007). In light of these results, it has been suggested that a gp350 vaccine could be improved upon with the inclusion of additional viral proteins (Cohen et al., 2013). Alternatively, it is possible that a vaccine targeting non-gp350 viral proteins could be more efficacious.

gH/gL is a promising antigen for vaccine development. Anti-gH/gL antibodies account for most serum antibodies that neutralize EBV infection of epithelial cells, but only a small fraction of antibodies that neutralize infection of B cells (Bu *et al*., 2019). Only a handful of anti-gH/gL monoclonal antibodies (mAbs) have been identified, all of which neutralize EBV infection of epithelial cells with comparable potency, but most have weak, or no neutralizing activity against EBV infection of B cells (Chesnokova and Hutt-Fletcher, 2011; Li et al., 1995; Molesworth *et al*., 2000; Sathiyamoorthy et al., 2016; Sathiyamoorthy et al., 2017; Snijder et al., 2018; Zhu et al., 2021). We previously described the isolation and characterization AMMO1, an anti-gH/gL monoclonal antibody (mAb) which potently neutralizes EBV infection of epithelial cells and B cells *in vitro* by binding to a discontinuous epitope on gH/gL (Snijder *et al*., 2018). The 769B10 mAb also neutralizes EBV infection of both cell types and binds to an epitope that overlaps with AMMO1, confirming this is a critical site of vulnerability on EBV (Bu *et al*., 2019). Passive transfer of AMMO1 severely limits viral infection following high-dose experimental EBV challenge in humanized mice and protects rhesus macaques against oral challenge with RhLCV if present at adequate levels at the time of challenge (Singh et al., 2020; Zhu *et al*., 2021). These studies provide proof of concept that anti-gH/gL antibodies can protect against EBV infection and indicate that a gH/gL-based vaccine capable of eliciting AMMO1-like antibodies could prevent oral transmission of the virus.

Here we generated several protein subunit vaccines where gH/gL is scaffolded onto self-assembling multimerization domains to produce nanoparticles with well-defined geometries and valency. Relative to monomeric gH/gL, immunization with the gH/gL nanoparticles elicited higher binding titers and neutralizing titers after one or two immunizations in mice. Competitive binding and depletion of plasma antibodies with an epitope-specific gH/gL probe suggested that only a small fraction of vaccine-elicited antibodies targeted the AMMO1 epitope. Consistent with this, depletion of plasma antibodies with an epitope-specific gH/gL knockout reduced plasma neutralizing activity to undetectable levels. Passive transfer of IgG purified from animals immunized with a computationally designed nanoparticle displaying 60 copies of gH/gL protected against high-dose lethal challenge in a humanized mouse model, while IgG purified from animals immunized with monomeric gH/gL did not. Collectively these results demonstrate that gH/gL is an attractive vaccine antigen, but that multivalent display of gH/gL is required to elicit neutralizing antibodies of sufficient titer to protect against EBV infection.

## Results

### Generation and Characterization of Multimeric gH/gL Vaccine Constructs

Cui *et al*. and Bu *et al*. have shown that immunization with multimeric gH/gL elicits higher serum neutralizing titers against infection of B cells and epithelial cells than immunization with monomeric gH/gL (Bu *et al*., 2019; Cui et al., 2016). However, these studies focused on a single multimerization platform when generating gH/gL constructs, either *Helicobacter pylori* ferritin, a 24-mer, or a T4 fibritin foldon domain, a trimer. Here we sought to develop several self-assembling multimeric gH/gL constructs with differing valencies, sizes, and geometries to evaluate how they differ in their ability to elicit neutralizing antibodies in mice. We generated various expression constructs where different multimerization domains were genetically fused to the C terminus of the gH ectodomain. These included i) a computationally designed circular tandem repeat protein (cTRP) that forms a planar toroid displaying four copies of gH/gL that is stabilized by inter-protomer disulfide bonds (Correnti et al., 2020); ii) a modified version of the multimerization domain from the C4b-binding protein from *Gallus gallus* (IMX313) which also forms a planar, ring-like structure stabilized by inter-protomer disulfide bonds capable of displaying seven copies of gH/gL (Ogun et al., 2008); iii) *Helicobacter pylori* ferritin which assembles into a 24-mer nanoparticle with octahedral symmetry and has previously been used to multimerize the EBV gp350 and gH/gL proteins (Bu *et al*., 2019; Kanekiyo et al., 2015); and iv) a secretion-optimized variant of a computationally designed, self-assembling 60-mer with icosahedral symmetry (Hsia et al., 2016). The gH fusion proteins were co-expressed with gL using the Daedalus lentiviral expression system in HEK293 cells (Bandaranayake et al., 2011). The gH/gL fusion proteins were purified by affinity chromatography followed by size-exclusion chromatography (SEC). The SEC elution profiles of the gH/gL fusion proteins were consistent with their expected valency (Fig. 1A). The 4-mer and 7-mer constructs eluted earlier than the monomer. The gH/gL 60-mer eluted in the void volume as expected, while the gH/gL 24-mer eluted near the void volume. Bands corresponding to the expected sizes of the gH fusion proteins were identified by SDS-PAGE (Fig. 1B). This analysis also revealed a band corresponding to gL and demonstrated that the preparations were highly pure (Fig. 1B).

**Figure 1.**
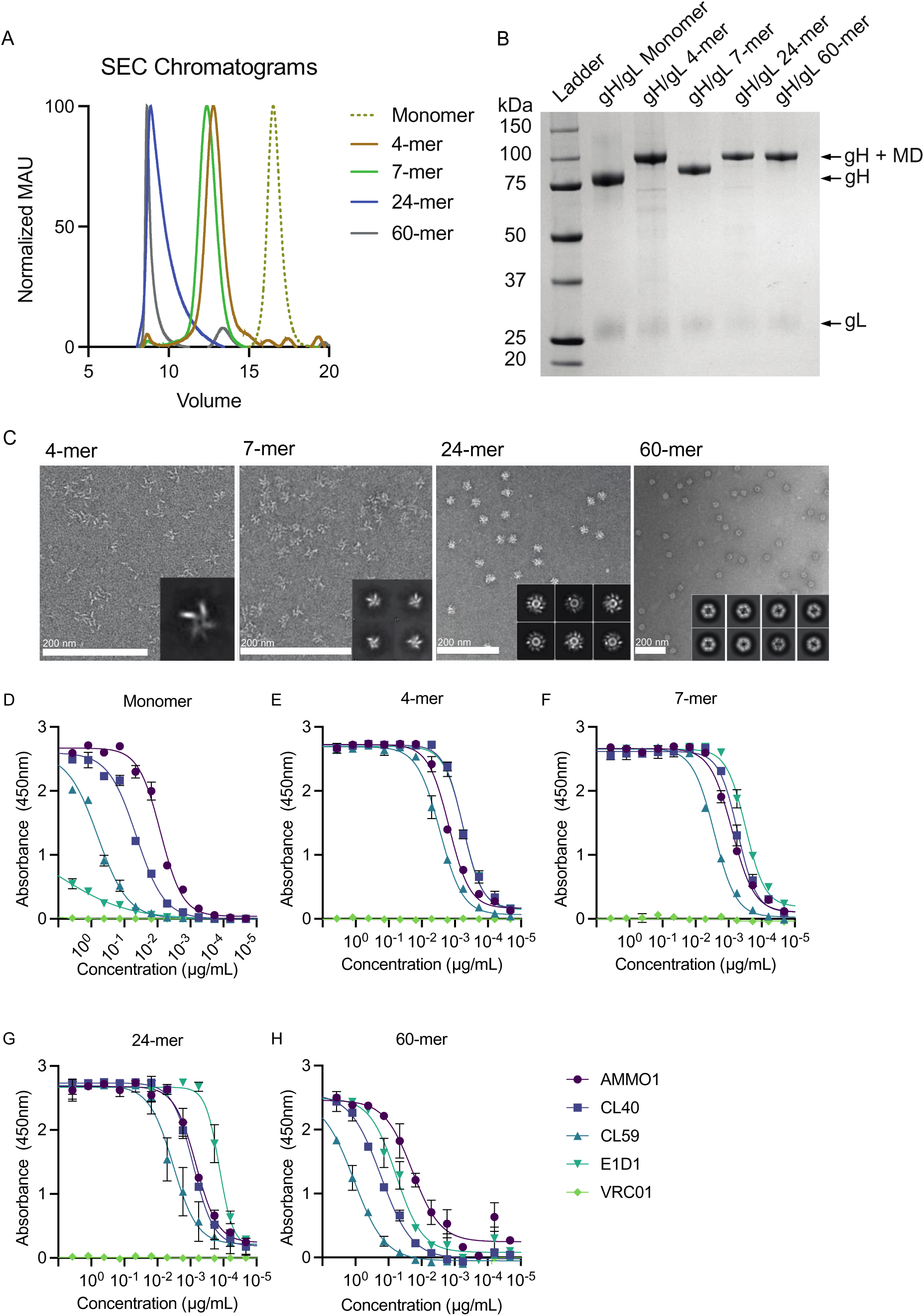
Biochemical and Biophysical Characterization of multimeric gH/gL nanoparticles. (**A**) Monomeric gH/gL and multimeric gH/gL nanoparticles were analyzed by size-exclusion chromatography (SEC) on a Superose 6 column as indicated. (**B**) Reducing SDS-PAGE analysis of 1 µg of monomeric gH/gL or multimeric gH/gL nanoparticles. Bands corresponding to gL, gH, and gH fused to 4-mer, 7-mer, 24-mer or 60-mer multimerization domains (MD) are indicated with arrows. (**C**) Negative stain electron microscopy was performed on 4-mer, 7-mer, 24-mer or 60-mer gH/gL nanoparticles as indicated. 2D class averages are shown in the inlay. Scale bars represent 200 nm. Binding of the anti-gH/gL mAbs E1D1, CL40, CL59 and AMMO1 to monomeric gH/gL **(D)** or multimeric gH/gL nanoparticles **(E-H)** were measured by ELISA as indicated. The anti-HIV-1 Env mAb VRC01 was used as a control for non-specific binding.

The gH/gL nanoparticles were imaged using negative-stain electron microscopy (nsEM), which demonstrated that all particles were monodisperse and of the predicted size. Density corresponding to gH/gL emanating from the nanoparticle cores was apparent in 2D class averages of the 4-mer, 7-mer and 24-mer (Fig. 1C). Density corresponding to gH/gL was less clearly defined on the 60-mer particles; however, a comparison of 3D reconstructions of the gH/gL-I3 fusions relative to the 60-meric I3 core revealed clear density corresponding to the C terminus of gH, indicating conformational flexibility around the gH-I3 fusion junction (Fig. S1).

To ensure that fusion to the multimerization domains did not alter the antigenicity of gH/gL, we measured the binding of several anti-gH/gL mAbs to each nanoparticle by ELISA. Of all the mAbs, AMMO1 binds with the highest affinity to monomeric gH/gL (Fig. 1D) (Snijder *et al*., 2018). The AMMO1 epitope bridges Domain I and Domain-II (D-I/D-II) and spans both gH and gL (Snijder *et al*., 2018). CL40 has the second highest affinity (Fig. 1D) (Snijder *et al*., 2018), and binds to an epitope spanning the D-II/D-III interface of gH (Sathiyamoorthy *et al*., 2017). CL59 binds at the C terminus of gH on D-IV (Sathiyamoorthy *et al*., 2017) and has lower affinity than CL40 or AMMO1 (Fig. 1D) (Snijder *et al*., 2018). E1D1 binds exclusively to gL and has the lowest affinity for the complex (Fig. 1D) (Sathiyamoorthy *et al*., 2016; Snijder *et al*., 2018).

In general, the mAbs maintained antigenicity to each multimeric construct, and some showed significant improvements in binding to the nanoparticles (Fig. 1D-H). Despite showing the weakest binding of all the mAbs to the gH/gL monomer, E1D1 showed the strongest binding to the 7-mer and the 24-mer (Fig. 1F and G). The E1D1 epitope is most distal to the multimerization domains and is therefore highly exposed on the nanoparticles. Moreover, the spacing of the E1D1 epitope may be optimally presented for bivalent engagement by the E1D1 mAb in some formats. In contrast, CL59 showed the weakest binding to all the gH/gL nanoparticles. CL59 binds closer to the C terminus of the gH ectodomain, which would be in close proximity to the nanoparticle core, potentially limiting exposure of the epitope in the context of the nanoparticles (Fig. 1E-H).

### Immunogenicity of gH/gL multimers

To assess the immunogenicity of the gH/gL nanoparticles, we immunized C57BL/6J mice with 5 µg of gH/gL monomer, 4-mer, 7-mer, 24-mer, or 60-mer formulated with adjuvant at weeks 0, 4 and 12. Plasma was collected two weeks post each immunization and endpoint binding titers to gH/gL were measured by ELISA (Fig. 2A). After the first immunization, the median reciprocal binding titers in the gH/gL 4-mer, 7-mer, 24-mer, and 60-mer groups were higher than those in the monomer group. A second immunization boosted the binding titers in each group 200- to 1000-fold. Again, the median titers in animals immunized with the gH/gL 4-mer, 7-mer, 24-mer, and 60-mer were higher than in those immunized with monomeric gH/gL.

**Figure 2.**
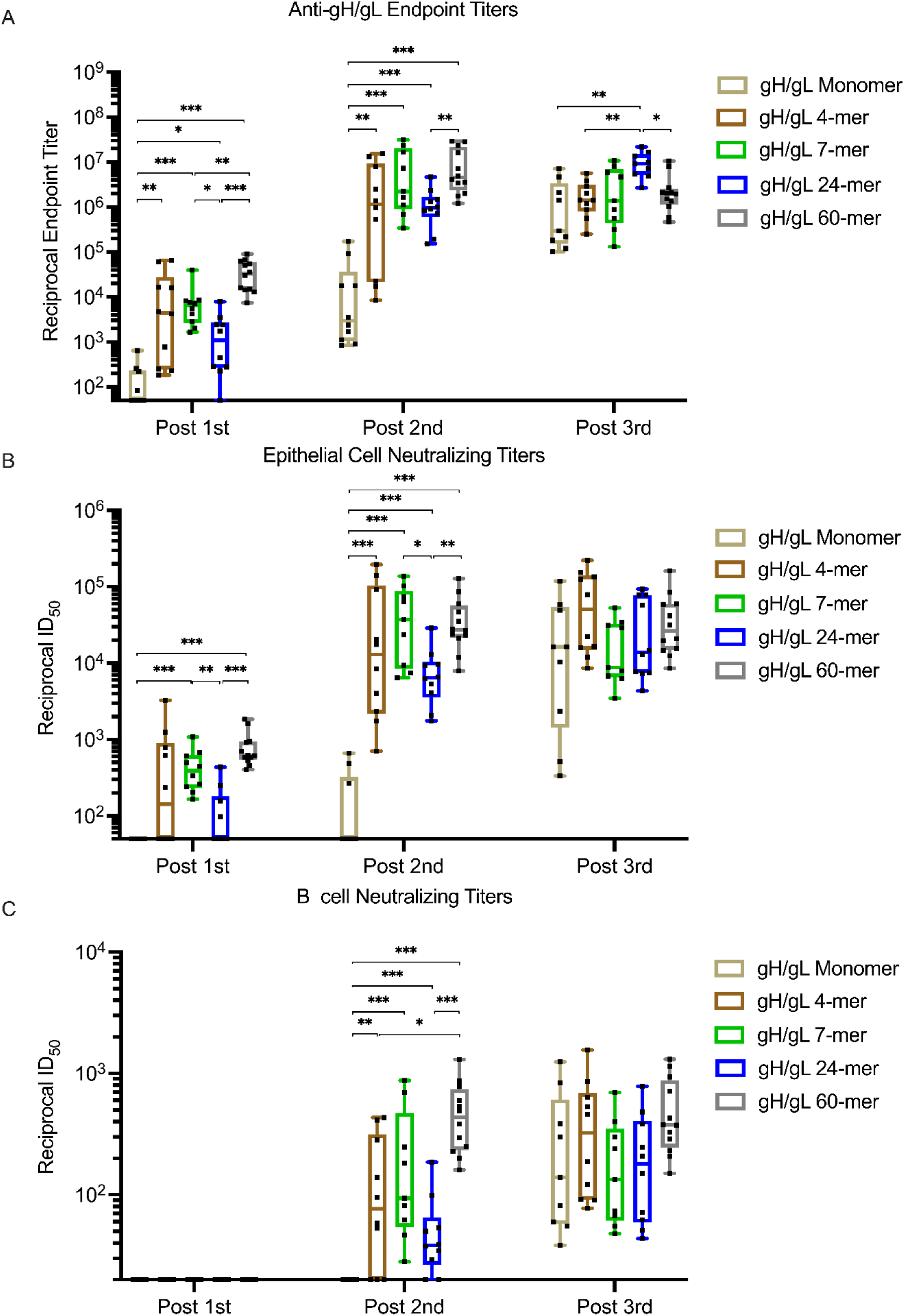
Immunogenicity of gH/gL nanoparticles. C57BL/6 mice were immunized with monomeric gH/gL or multimeric gH/gL nanoparticles at weeks 0, 4, and 12. Blood was collected 2 weeks after each immunization. (**A**) Endpoint plasma binding titers to gH/gL were measured by ELISA. Each dot represents the reciprocal endpoint titer for an individual mouse measured in duplicate. Box and whisker plots represent the minimum, 25^th^ percentile, median, 75^th^ percentile, and maximum values. The ability of plasma from individual mice to neutralize EBV infection of epithelial cells (**B**), or B cells (**C**). Each dot represents the reciprocal half-maximal inhibitory dilution (ID_50_) titer of an individual mouse. Plasma that did not achieve 50% neutralization at the lowest dilution tested (1:20) was assigned a value of 10. Box and whisker plots represent the minimum, 25^th^ percentile, median, 75^th^ percentile, and maximum values. Significant differences were determined using Mann-Whitney tests with Holm-adjusted p-values (*p < 0.05, **p < 0.01, ***p < 0.001)

A third immunization with the monomer boosted the gH/gL binding titers such that they were comparable to those elicited by the 4-mer, 7-mer, and 60-mer. A third immunization with the 24-mer also boosted the titers such that they were higher than the monomer 4-mer and 60-mer groups, while the third immunization with the other nanoparticles did not further boost the median binding titers (Fig. 2A).

We next measured the ability of vaccine-elicited plasma to neutralize EBV infection of both B cells and epithelial cells. Neutralizing activity against epithelial cell infection was elicited two weeks after the first immunization in all groups that received multimeric, but not monomeric gH/gL. The median reciprocal half maximal inhibitory dilution (ID_50_) titers were significantly higher in the 60-mer group compared to the monomer and 24-mer groups (Fig. 2B). Additionally, median titers were significantly higher in the 7-mer group compared to the monomer and 24-mer groups.

The second immunization boosted median neutralizing titers by ∼10-100 fold in the epithelial cell infection assay. The median neutralizing titers were higher in all of the gH/gL nanoparticle immunized groups than they were in the monomer group (Fig. 2B). The epithelial cell neutralizing titers in the 7-mer and 60-mer were also higher than those elicited by the 24-mer. The third immunization with the gH/gL nanoparticles did not further boost epithelial cell neutralizing responses, while the third dose of monomeric gH/gL boosted titers to levels that were comparable to those in other groups.

None of the gH/gL antigens elicited antibodies that could neutralize B cell infection two weeks after the first immunization (Fig. 2C). Following the second immunization, neutralizing titers were present in plasma from all groups immunized with gH/gL nanoparticles, but not in animals immunized with the monomer. Among the nanoparticle-immunized mice, the B cell neutralizing titers elicited by the 60-mer were higher than the 4-mer and 24-mer at this time point. Although the median B cell neutralizing titers elicited by the 60-mer (reciprocal ID_50_ = 436) were higher the 7-mer (reciprocal ID_50_ = 94), the difference was not statistically significant.

As was observed with the epithelial cell neutralizing titers, a third immunization with the gH/gL nanoparticles did not further boost B cell neutralizing responses, while a third dose of monomeric gH/gL boosted titers to levels that were comparable to those in other groups. In general, the neutralizing titers were about 10-fold lower against B cell infection compared to epithelial cell infection in all groups. From these analyses we conclude that all gH/gL nanoparticles displayed superior immunogenicity compared to monomeric gH/gL after one or two immunizations and that a third immunization did not result in a significant titer boost.

### Plasma epitope mapping

Each multimeric gH/gL nanoparticle tested here has a unique valency and geometry which differentially affects the exposure of certain epitopes bound by neutralizing anti-gH/gL mAbs (Fig. 1D-H). To test whether the nanoparticle format skewed the epitope-specificity of vaccine-elicited antibodies from each construct, we assessed the ability of pooled immune plasma to compete with the E1D1, CL40, CL59 and AMMO1 mAbs for binding to gH/gL by ELISA (Fig. 3A-D).

**Figure 3.**
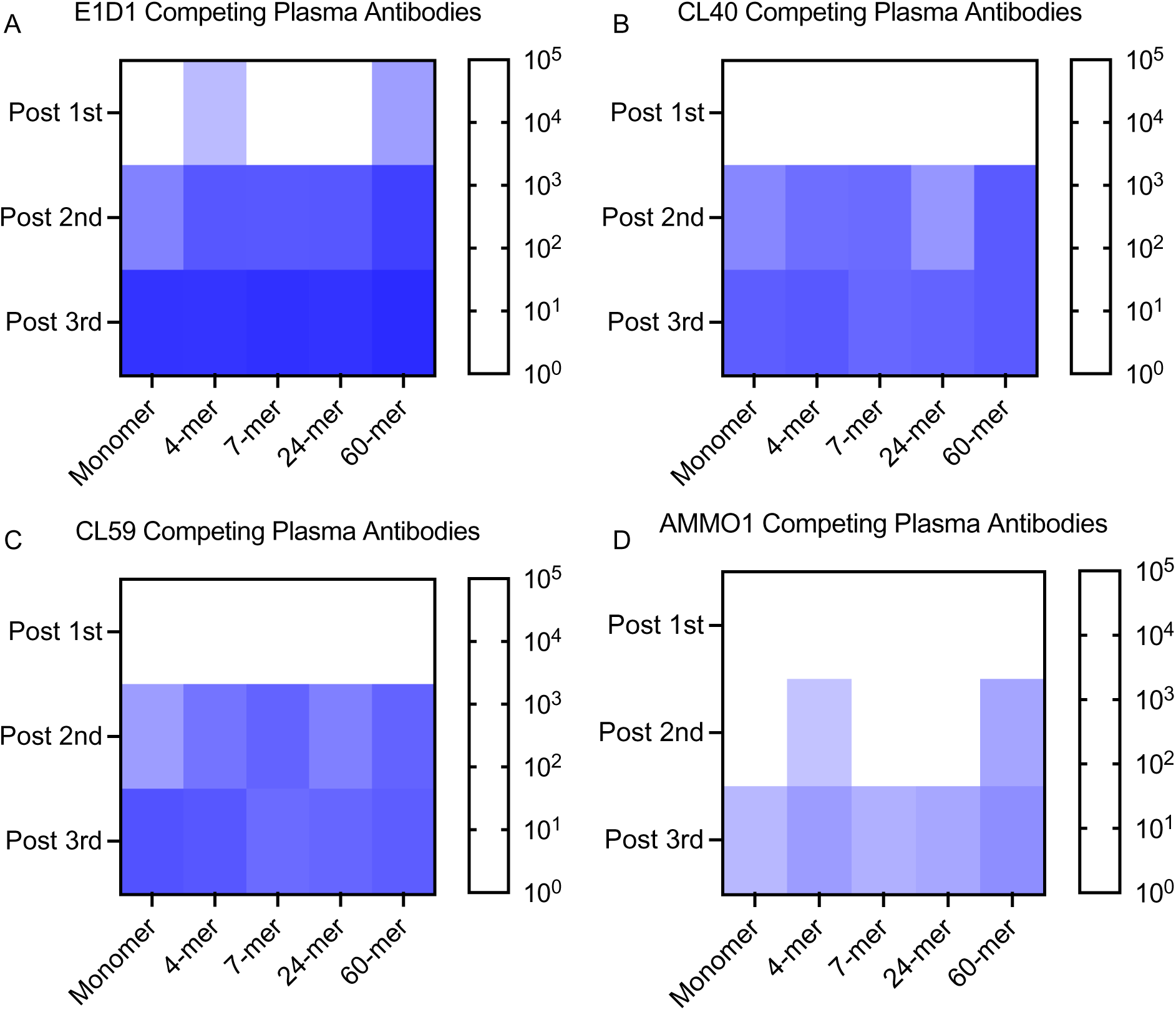
Plasma competition against monoclonal anti-gH/gL antibodies. The ability of plasma pooled from groups of mice immunized with monomeric gH/gL or multimeric gH/gL nanoparticles to inhibit binding to a panel of anti-gH/gL antibodies was measured by ELISA. The log reciprocal plasma dilution titers resulting in a 50% inhibition of (**A**) E1D1, (**B**) CL40, (**C**) CL59, or **(D**) AMMO1 antibodies at each time point, are shown on each heatmap as indicated. See Supplementary Figure S2 for titration curves.

Pooled plasma collected following one immunization with the gH/gL 4-mer and gH/gL 60-mer weakly inhibited E1D1 binding (Fig. 3A and S2A). After the second immunization, plasma from all groups inhibited CL40, CL59, and E1D1 binding (Fig. 3A-C and S2B). Plasma antibodies that inhibited binding of these mAbs were further boosted following a third immunization in most groups. The only exception was that a third immunization with the 7-mer did not boost CL59-blocking plasma antibodies (Fig. 3C and S2C).

Plasma antibodies capable of inhibiting AMMO1 binding were less common. Immune plasmas from the 4-mer and 60-mer groups weakly inhibited AMMO1 binding after two immunizations, and were boosted following a third immunization (Fig. 3D and S2C). All antigens elicited low titers of AMMO1-blocking antibodies following three immunizations. Among these, titers elicited by the 60-mer were highest at ∼1:150.

These experiments demonstrate that each gH/gL nanoparticle readily elicits antibodies that compete with E1D1, and that AMMO1-competing antibodies are rarer. This difference in competition could be attributed to the relative affinities of these mAbs for gH/gL (Fig. 1D), or it could be due to the relative exposure of these epitopes on the nanoparticle.

Although the titers of AMMO1-competing antibodies in the plasma of mice immunized with gH/gL nanoparticles are low, because the epitope boudn by this mAb represents a critical site of vulnerability on gH/gL, we sought to assess the relative contribution of AMMO1-like antibodies to the plasma neutralizing activity of immunized mice. To achieve this, we developed an epitope-specific gH/gL probe and carried out plasma depletions. We previously identified two mutations, K73W and Y76A that reduced binding of AMMO1 to cell-surface expressed gH/gL (Snijder *et al*., 2018). We expressed and purified a monomeric gH/gL ectodomain harboring these two mutations (herein called gH/gL-KO) which completely ablated AMMO1 binding while maintaining binding to other gH/gL mAbs as measured by biolayer interferometry (BLI) (Fig. 4 A-D).

**Figure 4.**
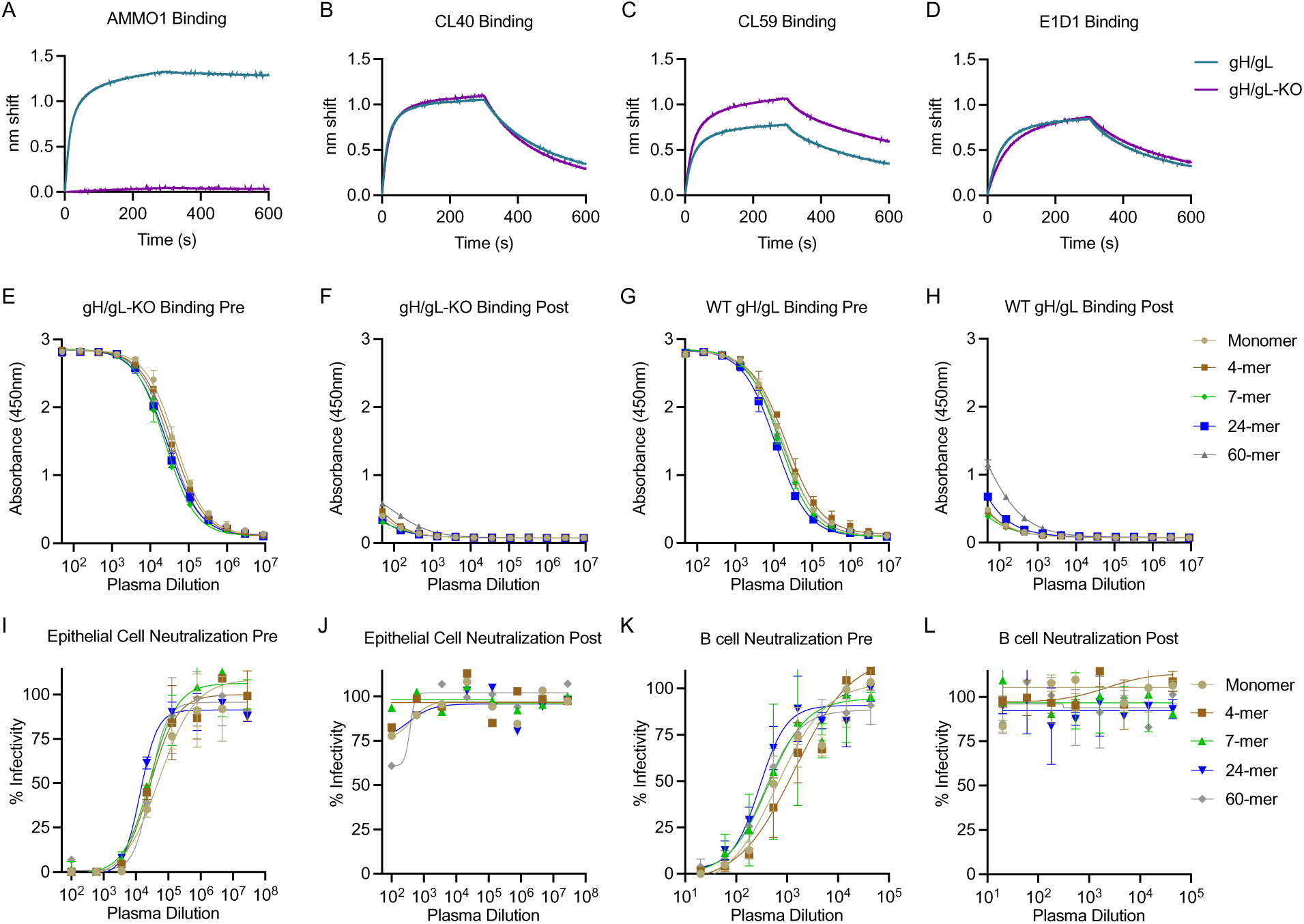
Depletion of non-AMMO1 like antibodies from pooled plasma. The binding of AMMO1 (**A**), CL40 (**B**), CL59 (**C**), and E1D1 (**D**) binding to gH/gL and gH/gL-KO (gH K73W,Y76A/gL) were measured using biolayer interferometry. **(E-H)** Antibodies were depleted from pooled plasma collected following three immunizations with gH/gL or gH/gL nanoparticles using gH/gL-KO conjugated magnetic beads. Pre- and post-depletion plasma samples were then assayed for binding to gH/gL and gH/gL-KO by ELISA as indicated. **(I-L)** The ability of plasma pre- and post-depletion to neutralize EBV infection was measured in B cells and epithelial cells.

Antibodies from pooled plasma collected from each group two weeks after the third immunization were depleted using immobilized gH/gL-KO. ELISA binding of depleted plasma to gH/gL-KO confirmed depletion of gH/gL-KO-specific antibodies (Fig. 4, compare panels E and F). Depletion with gH/gL-KO also reduced binding to wild-type gH/gL (Fig. 4G and H). The binding signal was slightly stronger for gH/gL relative to gH/gL-KO post-depletion (Fig. 4H and F), consistent with the mAb competition studies which demonstrated that there are very few AMMO1-like antibodies in the plasma of immunized animals.

Depletion of gH/gL-KO-specific antibodies led to a complete loss of neutralizing titers in both the B cell and epithelial neutralization assays (Fig. 4I-L). Collectively these data demonstrate that only a small portion of vaccine-elicited antibodies in each group target the AMMO1 epitope, and that they do not make a measurable contribution to the plasma neutralizing activity.

### Passive transfer of nanoparticle-elicited gH/gL mAbs protects against lethal challenge in humanized mice

Immunocompromised mice engrafted with human hematopoietic stem cells develop human B cells that can become infected by EBV and are used as an *in vivo* model of EBV infection (Fujiwara and Nakamura, 2020; Münz, 2017). This model has been used to evaluate the ability of monoclonal, or polyclonal antibodies elicited by either vaccination or infection to protect against controlled viral challenge (Cui et al., 2021; Kim et al., 2021; Singh *et al*., 2020; Zhu *et al*., 2021). Having established that gH/gL nanoparticles display superior immunogenicity, we sought to assess whether the antibodies they elicit confer protection against EBV challenge in this model.

To generate mice for these studies, non-obese diabetic [NOD] Rag1-/-, Il2rg-/- mice were engrafted with mobilized huCD34+ hematopoietic stem cells, hereafter referred to as humanized mice. At 20 weeks post-transplant, ∼10-25% of peripheral blood mononuclear cells were human cells, of which ∼65-80% were huCD19+ B cells (Fig. S3). Since the humanized mouse model used here does not efficiently generate antibody responses to immunization (Yu et al., 2017), we opted to passively transfer purified antibodies elicited in wild-type C57BL/6 mice, which allowed us to directly evaluate the protective efficacy of the vaccine-elicited antibodies independent of other vaccine-induced immune responses.

To generate sufficient antibody for these experiments, C57BL/6J mice (n=20) were immunized two times with 5 µg of the gH/gL 60-mer at weeks 0 and 4. This nanoparticle was selected because after two doses it consistently elicited high titers of antibodies that neutralize EBV infection of B cells which are the primary targets of infection in humanized mice. As a comparator, we also immunized a group of mice with gH/gL monomer (n=20). Two weeks after the second immunization, plasma were harvested, pooled, and total IgG was purified using protein A/G resin. As a control IgG was purifed from unimmunized C57BL/6J mice (n=20). The concentration of each pool of purified IgG was measured by spectrophotometry and gH/gL binding activity was measured by ELISA, demonstrating that the anti-gH/gL titers were higher in the 60-mer immunized group (Fig. S4).

IgG from each group were delivered to humanized mice (n=4-5 mice/group) via intraperitoneal injection 2 days prior to challenge at a dose of 500 µg IgG/mouse. Total IgG measured in pooled plasma collected 2 days prior to and 1 day after transfer confirmed that that the mice received comparable levels of total IgG (Fig. 5A). However, the levels of anti-gH/gL antibodies were higher in the mice that received IgG from 60-mer immunized animals compared to those that received plasma from animals immunized with the monomer (Fig. 5B), consistent with the superior immunogenicity of the gH/gL nanoparticle (Fig. 2A). Plasma from animals that received IgG from unimmunized animals (control IgG) did not display any binding activity to gH/gL (Fig. 5B). Two days after IgG transfer (day 0), each mouse was challenged via retro-orbital injection with 33,000 Raji Infectious Units (RIU) of EBV. We also included an infected control group which did not receive antibody pre-treatment and an uninfected control group that received neither antibody pre-treatment nor EBV challenge.

**Figure 5.**
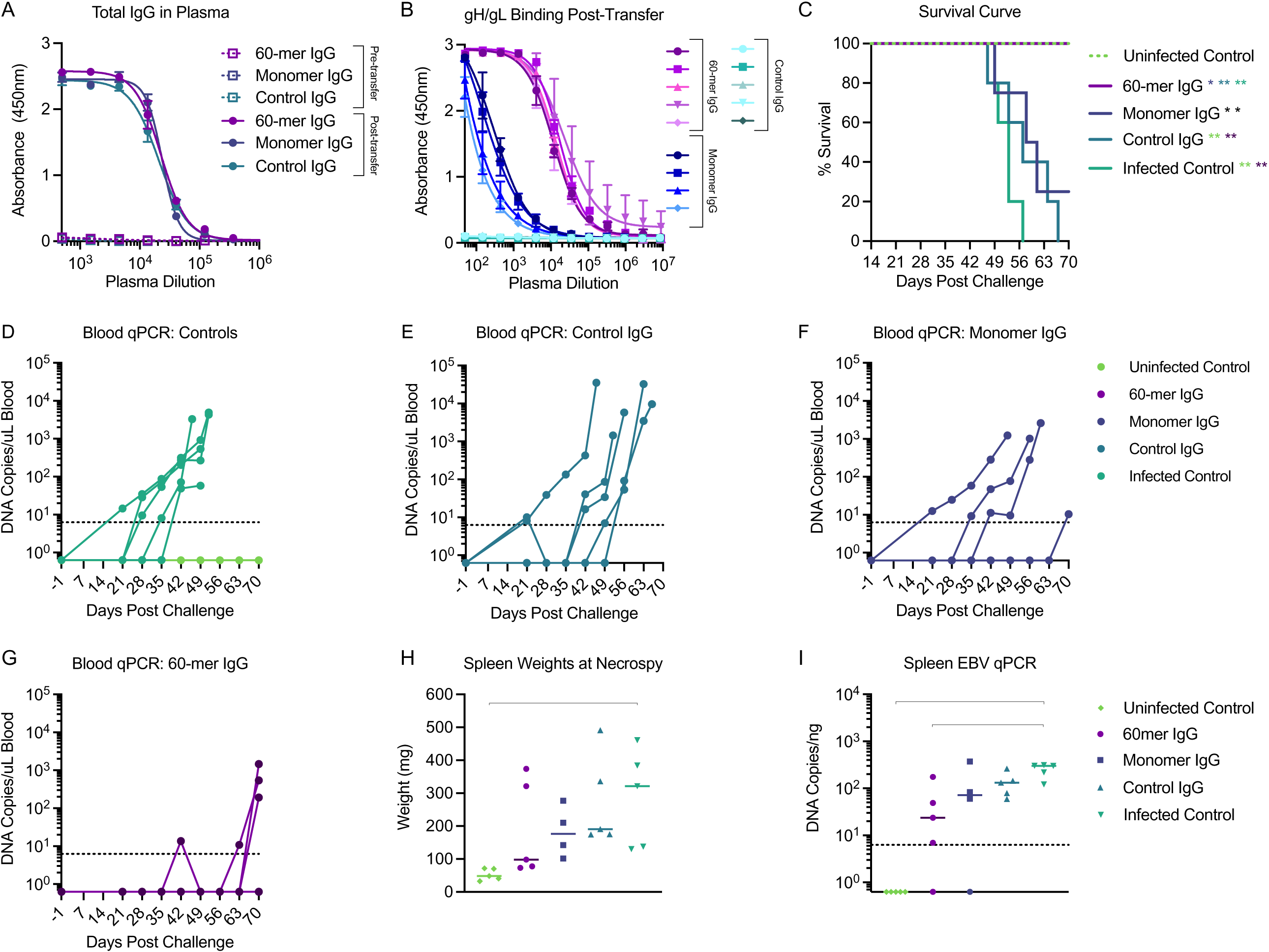
gH/gL nanoparticle elicited antibodies protect humanized mice from lethal EBV challenge. C57 BL6 mice were immunized with either monomeric or gH/gL 60-mer (n=10 per group) at weeks 0, and 4. Blood was collected by cardiac puncture at week 6, pooled and the serum IgG was purified. 0.5 mg of total IgG from monomer (n=4) or 60-mer (n=5) immunized mice was administered to humanized mice. A control group of mice received 0.5 mg total IgG purified from naïve C57 BL6 mice (n=5). (**A**) Total IgG was measured in pooled plasma from each group collected 3 days prior to and 1 day after IgG transfer. (**B**) Anti-gH/gL IgG antibodies from plasma collected from individual humanized mice one day after transfer was measured by ELISA as indicated. (**C**) Survival of humanized mice that received IgG purified from the indicated groups was monitored over a 70-day period following EBV challenge. An infected control group (n=5) did not receive IgG prior to challenge, and an uninfected control group (n=5) did not receive IgG or viral challenge. Significant differences in the survival data were determined using log-rank tests (*p < 0.05, **p < 0.01). **(D-G)** Viral DNA was quantified in the peripheral blood of infected and uninfected control (**D**), Control IgG (**E**), Monomer IgG (**F**), and 60-mer IgG (**G**) groups collected at the indicated timepoints via qPCR. Each series of connected dots represents an individual mouse at each time point analyzed, and the dashed line represents the limit of detection. (**H**) At necropsy, spleens were harvested and weighed. Each dot represents an individual mouse, and the bar represents the median weight in milligrams. Photographs of individual spleens are shown in Figure S6. (**I**) Viral DNA copy number was quantified in splenic DNA extracts at necropsy. Each dot represents an individual mouse, the bar represents the median copy number, and the dashed line indicates the limit of detection. Significant differences for spleen weight and viral DNA copy number were determined using Mann-Whitney tests with Holm-adjusted p-values (*p < 0.05).

Following challenge, the mice were weighed three times a week and monitored for general health over the course of 10 weeks. Blood samples were collected weekly beginning at 3 weeks post-challenge, and spleens were harvested from each mouse at the day 70 endpoint, or earlier if they met euthanasia criteria. Upon completion of the study, 100% of the animals in the uninfected control and 60-mer treatment groups survived (Fig. 5C). In contrast, 100% of the animals in the infected control and control IgG treatment groups succumbed to infection by 56- and 66-days post-challenge, respectively. 75% of mice in the monomer treatment group did not survive beyond day 60, and only one animal survived the entire 70 days (Fig. 5C). The survival rate of mice in the uninfected control group and 60-mer treatment group were significantly higher than all other groups (Fig. 5C).

EBV DNA was not detected in the blood or spleen of the uninfected control group throughout the duration of the experiment (Fig. 5D). In contrast, 100% of mice in the control IgG treatment group (Fig. 5E) and 100% of mice in the infected control group were viremic as early as 21 days post-challenge (Fig. 5D). In the monomer IgG group, EBV DNA was detected in the blood of 100% of mice (Fig. 5F). In the 60-mer IgG group, EBV DNA was undetectable in blood of 40% of mice at any time point tested. One mouse was viremic at weeks 7 and 10, another at weeks 9 and 10, and a third at week 10 (Fig. 5G).

A phenotypic analysis of the peripheral blood lymphocytes revealed a rapid decline in B cells in mice that received control IgG or IgG elicited by the monomer, and the infected controls relative to the uninfected controls approximately one-month post-challenge Fig. S3). A more gradual decline was noted in three of the mice that received IgG elicited by the gH/gL 60-mer (Fig. S3). This decline in B cell frequencies is consistent with T cell-mediated killing of infected B cells (Yajima et al., 2009).

At necropsy, spleens from animals in the infected control and control IgG groups were significantly heavier than those in the uninfected control groups (Fig. 5H) and had visible splenic tumors (Fig. S5). Spleens from two of the viremic mice in the 60-mer IgG group were about 3 times heavier than the other three mice in the 60-mer IgG group. All the spleens were enlarged in the monomer group relative to the uninfected control animals and had evidence of splenic tumors (Fig. 5H and S5). Viral DNA was not detected in the the spleens of all animals in the uninfected control group and from the spleens from one animal in each of the 60-mer IgG and monomer IgG groups (Fig. 5I), but it was detected in the spleens of all remaining mice (Fig. 5I).

Collectively, these data demonstrate that multivalent display of gH/gL elicits higher titers of neutralizing antibodies that protect against lethal EBV challenge in a humanized mouse model. However, they do not confer sterilizing immunity.

## Discussion

A safe and effective vaccine could alleviate the global disease burden resulting from EBV infection. Here we developed several multimeric vaccine candidates derived from the gH/gL ectodomain and evaluated their ability to elicit antibodies capable of neutralizing EBV infection of both B cells and epithelial cells in mice and demonstrated that a computationally designed nanoparticle displaying 60 copies of gH/gL elicited antibodies capable of protecting against high-dose, lethal challenge in a humanized mouse infection model.

Antigen multimerization has been used to improve the immunogenicity of subunit vaccines against several pathogens including malaria, HIV-1, RSV, SARS-CoV-2, influenza (Boyoglu-Barnum et al., 2021; Brouwer et al., 2019; Jardine et al., 2013; Jelínková et al., 2021; Kanekiyo et al., 2019; Marcandalli et al., 2019; Moon et al., 2012; Walls et al., 2021) and EBV (Bu *et al*., 2019; Cui *et al*., 2016; Cui *et al*., 2021). Multimerization can enhance the immunogenicity of subunit vaccines through several mechanisms including more efficient B cell receptor cross linking, triggering of innate B cell responses, lymph node trafficking, and enhanced MHC class II antigen presentation (Bachmann and Jennings, 2010; Irvine and Read, 2020; Irvine et al., 2013).

Previous studies have shown that antigen valency correlates with B cell activation, germinal center recruitment, and B cell differentiation as well as serum binding and neutralizing titers (Kato et al., 2020; Marcandalli *et al*., 2019; Veneziano et al., 2020). Although nanoparticles displaying gH/gL exhibited superior immunogenicity as compared to monomeric gH/gL, we did not observe a strict correlation between antigen valency and binding or neutralizing titers. The differences in the ability of these antigens to elicit neutralizing antibodies could be linked to nanoparticle stability *in vivo* or T cell help directed at MHCII-restricted epitopes that differ between the nanoparticle scaffolds (Arunachalam et al., 2021).

Because of its ability to potently neutralize infection of both cell types, the overlapping epitope targeted by AMMO1 and 769B10 represents a critical site of vulnerability on EBV (Bu *et al*., 2019; Snijder *et al*., 2018). Despite readily eliciting antibodies targeting several other epitopes on gH/gL, our analysis indicates that the AMMO1 epitope is subdominant in the context of immunization. Despite this, immune plasma from gH/gL 60-mer immunized mice was protective *in vivo*. Thus, relative to monomeric gH/gL, the gH/gL 60-mer may have elicited high titers of less potent anti-gH/gL antibodies like CL40 and CL59. Alternatively, the immunogens may have elicited antibodies targeting other potently neutralizing epitopes on gH/gL such as the one defined by the recently identified 1D8 mAb or other yet to be identified epitopes (Zhu *et al*., 2021). Gaining a better understanding of the epitopes on gH/gL that are targeted by neutralizing and non-neutralizing antibodies elicited by natural infection or immunization through the isolation and characterization of monoclonal antibodies would enable rational gH/gL vaccine design that could further enhance neutralizing titers when combined with multimeric antigen display. For example, immunogen design strategies could be employed to immunofocus the antibody response to potently neutralizing gH/gL epitopes.

The majority of humanized mice that received IgG elicited by monomeric gH/gL did not survive EBV challenge. Similarly, passive transfer of sera from rabbits immunized with monomeric gH/gL conferred partial protection from lethal EBV challenge in humanized mice (Cui *et al*., 2021). Since EBV infection in humanized mice is restricted to human B cells (Fujiwara and Nakamura, 2020; Münz, 2017), the observed lack of protection by IgG raised against monomeric gH/gL in our study is most likely due to the inability of this antigen to elicit antibodies that neutralize infection of B cells following two immunizations. In contrast, IgG purified from animals immunized with the gH/gL 60-mer prevented death following high dose EBV challenge, demonstrating that multivalent display substantially improves the quality of vaccine-elicited anti-gH/gL antibodies. Although we only evaluated the ability of antibodies elicited by the 60-mer in the humanized mouse challenge studies, the other nanoparticles developed here and elsewhere (Bu *et al*., 2019; Cui *et al*., 2021) have potential for clinical development and additional in vivo comparisons and manufacturing feasibility studies are warranted.

Both B cells and epithelial cells are present in the oropharynx, thus antibodies that can neutralize infection of both types of cells are an important consideration for EBV vaccine development (van Zyl et al., 2019). The gH/gL constructs evaluated here and elsewhere (Bu *et al*., 2019; Cui *et al*., 2021) consistently elicit higher epithelial cell neutralizing titers as compared to B cell neutralizing titers. Since murine epithelial cells are not susceptible to infection and oral transmission is not possible in humanized mice (Fujiwara et al., 2015; Münz, 2017), this challenge model may underestimate the relative importance of antibodies capable of neutralizing infection of this cell type. The evaluation of a multivalent gH/gL vaccine to prevent rhesus lymphocryptovirus infection of macaques, where oral transmission is the natural route of infection (Moghaddam et al., 1997), should more accurately predict its ability to protect humans against EBV.

In sum, we demonstrate that multivalent display of EBV gH/gL markedly enhanced immunogenicity in mice and that a computationally designed 60-mer nanoparticle elicited antibodies that protected against lethal challenge in a humanized mouse infection model. These results underscore the importance that vaccine-elicited antibodies against gH/gL can play in preventing EBV infection and highlight the utility of cutting-edge vaccine approaches in the development of vaccines against this important cancer-associated pathogen.

## Author Contributions

A.T.M and H.M. conceptuialized the study. H.M., L.J.H., Y-H.W, B.P., B.F., A.J.B., J.Y.Y., C.W., S.S., A.W., S.H., performed experiments and analyzed data. C.C., J.P., M.P., N.P.K., analyzed data. A.T.M. and H.M. wrote the initial manuscript, and all authors contributed to editing and figure preparation. Z.M. conducted statistical analysis. B.L.S., J.O., N.P.K., and A.T.M. Provided funding.

## Material and Methods

### Materials Availability

All materials generated herein are available upon request under an MTA from the corresponding author. The pTT3 vectors are used under license from the National Research Council of Canada. The UCOE element in the pCVL-UCOE0.7-SFFV based vectors is used under license from Millipore.

### Cell Lines

All cell lines were incubated at 37°C in the presence of 5% CO2 and were not tested for mycoplasma contamination. Raji cells (human male) were maintained in RMPI + 10% FBS, 2 mM L-glutamine, 100 U/ml penicillin, and 100 μg/ml streptomycin (cRPMI). 293-2089 cells (human female) were grown in cRPMI containing 100 μg/ml hygromycin (Delecluse et al., 1998). AKATA (human female) B cells harboring EBV in which the thymidine kinase gene has been replaced with a neomycin and GFP cassette virus (AKATA-GFP) were grown in cRPMI containing 350 μg/ml G418 (Molesworth *et al*., 2000). SVKCR2 cells (human male) were grown in DMEM containing 10% cosmic calf serum, 2 mM L-glutamine, 100 U/ml penicillin, 100 μg/ml streptomycin, 10 ng/ml cholera toxin and 400 μg/ml G418 (Li et al., 1992). 293-6E (human female, RRID:CVCL_HF20) and 293T cells (human female, RRID:CVCL_0063) cells were maintained in Freestyle 293 media with gentle shaking.

### Plasmids

pTT3 plasmids containing cDNA encoding gH (AA 19-679, Genbank AFY97969.1) with a C-terminal His and Avi tag, and gL AA 24-137 Genbank: AFY97944.1 with a TPA leader peptide have been previously described (Snijder *et al*., 2018). Site directed mutagenesis was used to introduce stop codons into gH between the His and Avi tags to produce an expression plasmid without the Avi tag, or 5’ of the His tag to produce an expression plasmid with no tags. The K73W and Y76A mutations to gH (herein called gH/gL-KO), were introduced into pTT3-gH-His using the QuickChange XL II Kit (Agilent, Cat. number #200521) according to manufacturer’s instructions.

To create a 7-mer gH expression construct, cDNA encoding a modified version of the C4b-BP protein (IMX313) (Ogun *et al*., 2008) followed by a stop codon was synthesized and cloned in-frame with the gH ectodomain in pTT3-gH-His-Avi, replacing the His and Avi tags to create pTT3-gH-IMX313. To create a 24-meric gH expression construct, cDNA encoding the gH ectodomain was amplified by PCR with primers that introduced an EcoRI site at the 5’ end followed by a (G_4_S)_2_ linker and finally a BamHI site at the 3’ end. The PCR amplicon was cloned into pTT3-426cTM4ΔV1-3-ferritin (Mcguire et al., 2016) (a kind gift from Dr. Leonidas Stamatatos) replacing the HIV-1 Env gene fused to *Helicobacter pylori* ferritin (Kanekiyo *et al*., 2015) to create pTT3-gH-ferritin.

cDNA encoding gH-IMX313 and gH-ferritin were amplified by PCR, and then cloned into the XhoI and BamHI restriction sites of pCVL-UCOE0.7-SFFV-muScn-IRES-GFP (Bandaranayake *et al*., 2011) replacing the muSCN cDNA, to create pCVL-UCOE0.7-SFFV-gH-IMX313-IRES-GFP and pCVL-UCOE0.7-SFFV-gH-ferritin-IRES-GFP. A C153T mutation which replaces an unpaired cysteine in gH was added to pCVL-UCOE0.7-SFFV-gH-IMX313-IRES-GFP using the QuickChange XL II Kit according to manufacturer’s instructions.

pCVL-UCOE0.7-SFFV-gH-I3-IRES-GFP was created by synthesizing a g-block encoding a modified version of I3-01 ((Hsia *et al*., 2016) and J.Y.W. manuscript in preparation) with homology to the 3’ end of the gH ectodomain at the 5’ end of the g block and homology to the downstream IRES region at the 3’ end of the g block. The plasmid backbone was amplified from pCVL-UCOE0.7-SFFV-gH-IMX313-IRES-GFP using a reverse primer that annealed to the 3’ end of the gH cDNA (containing the C153T mutation) and a forward primer that annealed to the 5’ end of the IRES and Platinum SuperFi II DNA polymerase (Thermo Fisher, catalog #12361010). The g block and linearized plasmid backbone were ligated together using the In-fusion HD cloning kit (Takara Bio, catalog #638920).

The tetramerization domain from cTRP24_6_SS (Correnti *et al*., 2020) was amplified by PCR using primers that added homology to the 3’ end of the gH ectodomain at the 5’ end homology to the downstream IRES region at the 3’ of the amplicon. The amplicon was ligated to the PCR linearized plasmid backbone described above using the In-fusion HD cloning kit to create from pCVL-UCOE0.7-SFFV-gH-cTRP(6)ss-IRES-GFP.

cDNA encoding gL was amplified by PCR, and then cloned into the XhoI and BamHI restriction sites of pCVL-UCOE0.7-SFFV-muScn-IRES-RFP (Bandaranayake *et al*., 2011) replacing the muSCN cDNA, to create pCVL-UCOE0.7-SFFV-gL-IRES-RFP. The sequences of all plasmids were confirmed by Sanger sequencing.

### Lentiviral Production

5.46 µg of psPAX2 (Addgene plasmid #12260), 2.73 µg of pMD2.G (Addgene plasmid #12259, both gifts from Didier Trono), and 11.05 μg of each pCVL-derived gH plasmid were mixed in 1.56mL PBS followed by 39µL of Freestyle Transfection Reagent (Millipore, catalog #72181). The transfection mix was gently agitated, incubated at room temperature for 15 minutes, and added dropwise to 13 mL of suspension-adapted 293T cells at 2×10^6 cells/mL in a 125 mL flask. After 24 hours, an additional 15 mL of 293 Freestyle media containing 15 µg of valproic acid was added to the cell culture. After another 48 hours, the cell culture was centrifuged at 1000 × *g* for 3 minutes, the supernatant was passed through a 0.44 µm filter, aliquoted, and stored at -80°C.

### Lentiviral Transduction

Polybrene (Millipore Sigma, catalog # TR-1003-G) was added to 10 mL of 293E cells at 1×10^6^ cells/mL to a final concentration of 2 µg/mL in addition to 2-3 mL of supernatant containing lentiviral particles harboring the various gH and gL expression constructs. 24 hours following transduction, 15 mL of 293Freestyle media (ThermoFisher, catalog #12338018) was added to the culture. A Guava easyCyte Flow Cytometer was used to monitor gH (GFP+) and gL (RFP+) transduction efficiency 72 hours after transduction. Transduced cultures were expanded to a total volume of 1 liter and cultured until cell viability declined to ∼80%. The transduced cell cultures were centrifuged at 4000 × *g* for 10 minutes to pellet cells. The supernatant was further clarified by passing through a 0.22 µm filter.

### Purification of untagged monomeric gH/gL

Clarified cell supernatant was adjusted to pH 5.5-6 using 2 M acetic acid. The clarified cell supernatant was incubated with CaptoMMC resin, pre-equilibrated with 30 mM sodium acetate, 50 mM NaCl, pH 5.5 (MMC binding buffer), then washed with 10 column volumes of MMC binding buffer and then eluted with 10 column volumes of 50 mM sodium acetate, 1 M ammonium chloride, pH 7.4. The protein elute was collected and concentrated using an Amicon Ultra-4 Centrifugal Filter Unit (Millipore Sigma, UFC803024), and further purified via size exclusion chromatography (SEC) on a HiLoad 16/600 Superdex 200 pg column (Millipore Sigma, 28-9893-35) with 10 mM Tris, 50 mM NaCl, pH 7.4 as the mobile phase. The protein was further purified by anion exchange chromatography using a HiTrap Q HP column (Cytivia, 29-0513-25) pre-equilibrated with 10 mM Tris, 50 mM NaCl, pH 7.4. The column was washed with 7% elution buffer (10 mM Tris, 1 M NaCl, pH 7.4) until the absorbance at A280 achieved a stable baseline. gH/gL was eluted over a linear gradient from 7% to 25% elution buffer over 20 column volumes. The eluted protein was further purified by SEC with Phosphate-Buffered Saline (PBS, 1×, Corning, 21-040-CM) as the mobile phase on the Superose 6 Increase 10/300 GL (Millipore Sigma, 29-0915-96). Fractions were analyzed by SDS-PAGE to identify those containing gH/gL >95% purity based on Coomassie blue staining. The purified protein was aliquoted, flash frozen in liquid nitrogen and stored long term at -80°C.

### Purification of Polyhistidine Tagged Proteins gH/gL His, gH/gL-KO, and gH/gL-cTRP(6)ss

Clarified cell supernatant was adjusted to a final concentration of 10 mM imidazole and 500 mM NaCl and then incubated with Ni-NTA resin (ThermoFisher, 88221) pre-equilibrated with 10 mM Tris, 500 mM NaCl, 10 mM imidazole, 0.02% azide, pH 7.1 (Ni-NTA binding buffer). The column was then washed with 10 column volumes of Ni-NTA binding buffer and eluted using 10 mM Tris, 500 mM NaCl, 500 mM imidazole, pH 8.0. The NiNTA eluate was subsequently purified by SEC using a Superdex 200 column with PBS as the mobile phase. Purified protein was aliquoted flash frozen in liquid nitrogen and stored at -80°C.

### Purification of gH/gL-4-mer (gH/gL-IMX313) and gH/gL-ferritin

Clarified cell supernatant was adjusted to a final concentration of 100 mM NaCl and then incubated with Galanthus Nivalis Lectin Agarose (Vector Laboratories, AL-1243-5), washed with 10 column volumes of 20 mM Tris, 100 mM NaCl, 1 mM EDTA, pH 7.4 and eluted with 20mM Tris, 100 mM NaCl, 1 mM EDTA, 1M methylmannopyranoside, pH 7.4. The eluted protein was further purified by SEC with PBS as the mobile phase on the Superdex 200 column or the Superose 6 Increase 10/300 GL for gH/gL-C4b and gH/gL-ferritin, respectively. gH/gL-IMX313 was flash frozen and stored at -80°C. gH/gL-ferritin was expressed and purified within a week of each immunization and stored at 4°C.

### Purification of gH/gL-I3

To prepare an affinity chromatography resin to purify gH/gL-I3, 4.5 mg E1D1 antibody was incubated with 1 mL Protein A resin (GoldBio, P-400) with rotation at room temperature for 30 minutes and then washed thoroughly with PBS. 6.5 mg of disuccinimidyl suberate (Pierce, 21555) was dissolved in 0.5 mL DMSO, then diluted in 10 mL PBS and added to the Protein A resin. The resin and DSS mixture were incubated at room temperature with rotation for at least 1 hour. The resin was washed thoroughly with PBS, and then incubated overnight with rotation at 4°C in 10 mL of 1 M Tris, pH 7.5, and washed again extensively with PBS. The resin was then washed with Pierce IgG Elution buffer (ThermoFisher Scientific, catalog #21004) to remove any E1D1 antibody that was not crosslinked to the resin, and then washed again with PBS. The E1D1 affinity resin was stored in 50 mM Tris, 150 mM NaCl, 0.02% azide when not in use.

Supernatant from cells transduced with gH/gL-I3-was incubated with the E1D1 resin, washed with TBS, eluted with Pierce IgG elution buffer, and neutralized with a 1/10^th^ volume of 1 M Tris pH 8. The eluted protein was further purified by SEC using a Superose 6 Increase column with 50 mM Tris, 150 mM NaCl, 150 mM L-arginine, pH 8 as the mobile phase. Purified protein was flash frozen and stored at -80°C.

### Recombinant Antibodies

Cloning, expression and purification of AMMO1 (Snijder *et al*., 2018), CL40, and CL59 (Singh *et al*., 2020) was performed as previously described. For cloning of E1D1, codon-optimized cDNA corresponding to E1D1 VH (GenBank: KX755644) was synthesized (Integrated DNA Technologies) and cloned in-frame with the human IgG1 constant region in pTT3-based expression vectors. Codon-optimized cDNA corresponding to E1D1 VL (GenBank: KX755645) was cloned in-frame with the human kappa constant regions in pTT3-based expression vectors. Recombinant E1D1 was expressed in 293-E cells and purified using Protein A affinity chromatography.

### Negative-Stain Electron Microscopy

For gH/gL 4-mer and 7-mer, 1% uranyl formate negative staining solution and Formvar/carbon grids (Electron Microscopy Sciences) of 300 mesh size were used to perform the negative staining experiment. The protein samples of 4-mer and 7-mer gH/gL were diluted to ∼40 μg/ml and ∼50 μg/ml, respectively and applied for 60 sec on glow discharged grids. Excess sample was blotted off using Whatman filter paper and the grids were rinsed using water droplets and further stained for additional 60 seconds. Excess stain was blotted off and the grids were air dried for 1-2 min.

For gH/gL 24-mer and 60-mer, sample were diluted to 100 µg/mL and 3 µL was negatively stained using Gilder Grids overlaid with a thin layer of carbon and 2% uranyl formate as previously described (Veesler et al., 2014).

For the gH/gL 4-mer and 7-mer, data were collected using a FEI Tecnai T12 electron microscope operating at 120 keV equipped with a Gatan Ultrascan 4K×4K CCD camera. The images were collected using an electron dose of 45.05 e−/Å^2^, a magnification of 67,000 × that corresponds to pixel size of 1.6 Å, and exposure time of 1 sec. The defocus range used was -1.00 µm to -2.00 µm. The data was collected using Leginon interface (Suloway et al., 2005) and processed using cisTEM (Grant et al., 2018). CTF correction on the collected micrographs was done using CTFFIND (Rohou and Grigorieff, 2015) within cisTEM. Particles were further picked from the micrographs and subjected to 2D classification and the best 2D classes were selected.

For the gH/gL 24-mer, data were collected on an FEI Technai 12 Spirit 120kV electron microscope equipped with a Gatan Ultrascan 4000 CCD camera. A total of 150 images were collected per sample by using a random defocus range of 1.1–2.0 µm with a total exposure of 45 e−/A^2^. Data were automatically acquired using Leginon, and data processing was carried out using Appion (Lander et al., 2009). The parameters of the contrast transfer function (CTF) were estimated using CTFFIND4 (Mindell and Grigorieff, 2003), and particles were picked in a reference-free manner using DoG picker (Voss et al., 2009). Particles were extracted with a binning factor of 2 after correcting for the effect of the CTF by flipping the phases of each micrograph with EMAN 1.9 (Ludtke et al., 1999). The gH/gL 24-mer stack was pre-processed in RELION/2.1 (Kimanius et al., 2016; Scheres, 2012a; b) with an additional binning factor of 2 applied, resulting in a final pixel size of 6.4 Å. Resulting particles were sorted by reference-free 2D classification over 25 iterations.

For the 60-mer, data were collected on an Talos L120C 120kV electron microscope equipped with a CETA camera. A total of ∼350 images were collected per sample by using a random defocus range of 1.3–2.3 μm, with a total exposure of 35 e−/A^2^, and a pixel size of 3.16 Å/pixel. Data were automatically acquired using EPU (ThermoFisher Scientific). All data processing was performed using CryoSPARC (Punjani et al., 2017). The parameters of the contrast transfer function (CTF) were estimated using CTFFIND4 (Mindell and Grigorieff, 2003), and particles were picked initially in a reference-free manner using blob picker, followed by template picking using well-defined 2D classes of intact nanoparticles. Particles were extracted after correcting for the effect of the CTF for each micrograph with a box size of 256 pixels. Extracted particles were sorted by reference-free 2D classification over 20 iterations. 3D ab initio was performed in cryoSPARC with the subsequent homogenous refinement step performed using icosahedral symmetry. The resulting 3D map was displayed at two different contours levels and images were generated using ChimeraX (Pettersen et al., 2021).

### Mice

Mice were housed in a specific pathogen-free facility at Fred Hutchinson Cancer Research Center. The facility is accredited by the Association for Assessment and Accreditation of Laboratory Animal Care. Mice were handled in accordance with the Guide for the Care and Use of Laboratory Animals (National Research Council of the National Academies), and experiments were approved by the Fred Hutch Institutional Animal Care and Use Committee and Institutional Review Board.

### Humanized Mice

NOD-scid Il2rg^null^ (NSG) NSG mice were irradiated with 275 Roentgen and then engrafted with 1×10^6^ CD34-enriched PBSCs obtained from granulocyte colony-stimulating factor mobilized healthy donors purchased from the Co-operative Center for Excellence in Hematology, Fred Hutchinson Cancer Research Center, by intravenous injection. 10 weeks post-cell transfer, successful human cell engraftment was confirmed via immunophenotyping of circulating lymphocytes using antibodies at indicated dilution: hCD45-FITC (1:100, eBioscience, catalog #11-9459-41), hCD8-BV21 (1:100, BD, catalog #562429), L/D-BV506 (1:200, Invitrogen, catalog #65-0866-14), hCD19-BV711 (1:100, Biolegend, catalog #302246), hCD20-BV786 (1:200, BD, catalog #743611), mCD45 (1:200, Biolegend, catalog #103112), hCD4-A700 (1:250, Invitrogen, catalog #56-0048-92), hCD33-PE (1:100, Biolegend, catalog #333404), mFcBlock (1:200, Biolegend, catalog #101302).

### Immunizations in C57BL/6 Mice

Comparative immunogenicity studies were performed in groups of 10 C57BL/6 mice (5 male and 5 female) between 7-10 weeks of age. After collecting a pre-bleed, mice were immunized at weeks 0, 4, and 12 with 5 µg (total protein) of monomer, 4-mer, 7-mer, 24-mer, or 60-mer formulated with 20% (v/v) synthetic lipid A in squalene emulsion SLA-SE (Carter et al., 2016) (purchased from the Infectious Disease Research Institute) in PBS or TBS (60-mer only) at a total volume of 100 µL. Mice were immunized via intramuscular injection split into two 50 µL doses split between both rear legs. Blood was collected retro-orbitally 2 weeks after the first and second immunizations and via cardiac puncture at week 14. Blood was collected in tubes containing a 1/10^th^ volume of citrate. Plasma was separated from whole blood via centrifugation and then heat inactivated at 56°C for 30 min. For passive transfer experiments into humanized mice immunizations were performed in groups of 20 C57BL/6 mice (10 male and 10 female) between 7-10 weeks of age. After collecting a pre-bleed, mice were immunized at weeks 0 and 4 with 5 µg of gH/gL monomer or 60-mer formulated in PBS (monomer) or TBS (60-mer) with 50% (v/v) Sigma Adjuvant System (SAS; Sigma Aldrich, catalog #S6322) for a total volume of 100 µL. Mice were immunized via intramuscular injection split 50 µL each between both rear legs. Blood was collected retro-orbitally via cardiac puncture at week 6 into a separate vial for each mouse containing 100 μL citrate. Plasma was separated from whole blood via centrifugation.

### IgG Purification from Murine Plasma

Plasma was pooled and heat inactivated at 56°C for 1 hour then diluted in protein G binding buffer (Pierce ThermoFisher, catalog #54200) and passed over a column containing 1mL of protein A/G resin (Pierce ThermoFisher, catalog #20422). The column was then washed 3 times with 5 column volumes of binding buffer. Finally, IgG was eluted from the resin in 5 × 2 mL fractions using IgG elution buffer (Pierce ThermoFisher, catalog #21004). Fractions were buffer exchanged into PBS, concentrated, filter sterilized, and yields were measured by nanodrop.

### EBV-Reporter Virus Production

To produce B-cell tropic GFP reporter viruses (B95-8/F), 9×10^6^ 293–2089 cells were seeded on a 15 cm tissue culture plate in cRPMI containing 100 μg/mL hygromycin. 24 hours later the cells were washed twice with PBS, the media was replaced with cRMPI without hygromycin, and the cells were transfected with 15 μg of each of p509 and p2670 expressing BZLF1 and BALF4, respectively, using GeneJuice transfection reagent (Sigma Aldrich, catalog #70967). 72 hours later the cell supernatant was collected, centrifuged at 300 × *g* for 5 min and then passed through a 0.8 μm filter. To produce epithelial cell tropic virus, B cells harboring AKATA-GFP EBV were suspended at 4×10^6^ cells/ml in RPMI containing 1% FBS. Goat anti-human IgG (SouthernBiotech, catalog #1030-01) was added to a final concentration 100 μg/ml and incubated at 37°C for 4 hours. Cells were then diluted to 2×10^6^ cells/ml in RPMI containing 1% FBS and incubated for 72 hours. Cells were pelleted by centrifugation at 300 × *g* for 10 min and then the supernatant was passed through a 0.8 μm filter. Bacitracin was added to a final concentration of 100 μg/mL. Virions were concentrated 25× by centrifugation at 25,000 × *g* for 2 hours and re-suspended in RPMI containing 100 μg/ml bacitracin. Virus was stored at -80°C and thawed immediately before use.

### Statistics

Kruskal-Wallis tests were performed to assess whether the distributions of responses varied across treatment groups, with p-values < 0.05 considered significant. If the Kruskal-Wallis test reached significance, a Mann-Whitney test was used to compare the distribution of outcomes between the pairs of groups considered. Immunogenicity was compared across each pair of treatment groups; for spleen weights and viral DNA copies, each group was compared to the infected control. The Holm method was used to adjust for multiplicity across the Mann-Whitney tests conducted for each outcome, with Holm’s adjusted p-values reported. For survival data, significant differences were determined using Log-rank Mantel-Cox test.

### B cell Neutralization Assay

B cell neutralization assays were carried out in Raji cells essentially as described (Sashihara *et al*., 2009). Mouse plasma was serially diluted in duplicate wells of 96 well round-bottom plates containing 25 µL of cRPMI. 12.5 µl of B95-8/F virus (diluted to achieve an infection frequency of 1-5% at the final dilution) was added to each well and plates were incubated at 37°C for 1 hour. 12.5 µl of cRPMI containing 4×10^6^ Raji cells/ml was added to each well and incubated for another hour at 37°C. The cells were then pelleted, washed once with cRPMI, and re-suspended in cRPMI. Reciprocal plasma dilutions are reported relative to the final infection volume (50 µL). After 3 days at 37°C, cells were fixed in 2% paraformaldehyde. The percentage of GFP+ Raji cells was determined on a BD LSRII cytometer or Luminex Guava HT cytometer.

To account for any false positive cells due to auto-fluorescence in the GFP channel, the average %GFP+ cells in negative control wells (n = 4-6) was subtracted from each well. The infectivity (%GFP+) for each well was plotted as a function of the plasma dilution. The neutralization curve was fit using the log(inhibitor) versus response-variable slope (four parameters) analysis in Prism 9.2.0 (GraphPad Software). The half maximal inhibitory plasma dilution ID_50_ was interpolated from the curve in Prism 9.2.0 (GraphPad Software).

For depletion assays, the average %GFP+ cells in negative control wells (n=4-6) was subtracted from each well. The %Infectivity was calculated for each well by dividing the %GFP+ cells in each well by the average %GFP+ cells in the most dilute plasma dilution wells and multiplying by 100. %Infectivity was plotted as a function of the plasma dilution. The neutralization curve was fit using the log(inhibitor) versus response-variable slope (four parameters) analysis in Prism 9.2.0 (GraphPad Software).

### Epithelial Cell Neutralization Assay

1.5 × 10^4^ SVKCR2 cells per well were seeded into a 96 well tissue culture plate. The following day plasma was serially diluted in duplicate wells containing 20 μL of media in a 96 well flat bottom plate followed by the addition of 20 μL of 25× concentrated epithelial cell-tropic virus and incubated for 15 min. Media was aspirated from the SVKCR2 cells and replaced by the antibody-virus mixture and incubated at 37°C. 48 hours later the cells were detached from the plate using 0.25% trypsin (Gibco, catalog #25200056), transferred to a 96 well round bottom plate, washed twice with PBS, and fixed with 10% formalin (Sigma Aldrich, catalog #HT501128), and the percentage of GFP+ cells were determined on an BD LSRII cytometer or Luminex Guava HT. Percent neutralization was determined as in the B cell neutralization assay.

### Measurement of Plasma Antibody Endpoint Binding Titers by Anti-His Capture ELISA

30µL/well of rabbit anti-His tag antibody (Sigma Aldrich, catalog #SAB5600227) was adsorbed at a concentration of 0.5 μg/mL on to 384 well microplates (Thermofisher, catalog #464718) at 4°C for 16 hours in a solution of 0.1 M NaHCO3 pH 9.4-9.6 (coating buffer). The next day, plates were washed 4 times with 1 x PBS, 0.02% Tween 20 (ELISA wash buffer) prior to blocking for 1 hour with 80 µL/well of 1× PBS containing 10% non-fat milk and 0.02% Tween 20 (blocking buffer). After blocking, plates were washed 4× with wash buffer and 30 µl/well of a 2 µg/mL solution of His-tagged gH/gL diluted in blocking buffer was added to the plate and incubated for 1 hour, and then washed 4× with ELISA wash buffer. Plasma was diluted in blocking buffer and three-fold serial dilutions were performed in duplicate followed by a 1-hour incubation at 37°C. 8-16 additional control wells were included that contained immobilized gH/gL but no immune plasma (control wells). Following 4 additional washes with ELISA wash buffer, a 1:2,000 dilution of goat anti-mouse IgG (SouthernBiotech, catalog #1033-05) in blocking buffer was added to each well and incubated at 37°C for 1 hour followed by 4 washes with ELISA wash buffer. 30 µl/well of SureBlue Reserve TMB Microwell Peroxidase substrate (SeraCare, catalog #5120-0081) was added. After 5 min, 30μL/well of 1N sulfuric acid was added and the A_450_ of each well was read on a Molecular Devices SpectraMax M2 plate reader. The binding threshold was defined as the average plus 10 times the standard deviation of the determined by calculating the average of A_450_ values of the control wells. Endpoint titers were interpolated from the point of the curve that intercepted the binding threshold using the Prism 9.2.0 package (GraphPad Software).

### Measure of Competitive Binding Titers by ELISA

Coating, blocking, and gH/gL immobilization steps were performed as described under “Measurement of plasma antibody endpoint binding titers by anti-His capture ELISA.” Following gH/gL capture, equal amounts of plasma from each mouse in a group were pooled and diluted in blocking buffer and 2-fold serial dilutions were performed, followed by a 1-hour incubation at 37°C. Following 4 additional washes with ELISA wash buffer, monoclonal antibodies AMMO1, CL40, CL59, and E1D1 were added at a concentration that achieves half-maximal binding (EC_50_; pre-determined in the same assay in the absence of competing sera) to each well containing serially diluted pooled sera from each group, followed by a 1-hour incubation at 37°C. After 4 washes with ELISA washing buffer, a 1:20,000 dilution of goat anti-human IgG (Jackson ImmunoResearch, catalog #115-035-008) in blocking buffer was added to each well and incubated at 37°C for 1 hour followed by 4 washes with ELISA wash buffer. Addition of SureBlue Reserve TMB Microwell Peroxidase substrate, addition of 1N sulfuric acid, and reading of plates was performed as described above. The average A_280_ values of buffer only control wells were subtracted from each mAb containing well and plotted in Prism 9.2.0 (GraphPad Software). A280 values were plotted as a function of the log_10_ of the plasma dilution. A binding curve was fit using the Sigmoidal, 4PL, X is log(concentration) least squares fit function. Maximum binding was defined as the best-fit value for the top of each curve computed in Prism. A_280_ values at each dilution on the curve were divided by the maximum binding and multiplied by 100 to calculate the % of max binding ([A_280_ at each dilution/max binding] × 100). The titer at which half-maximal binding was observed was interpolated from the binding curve using the Prism 9.2.0 package (GraphPad Software).

### Biotinylation of recombinant proteins

Recombinant gH/gL proteins were biotinylated using the EZ-Link NHS-PEG4-Biotin Kit (ThermoFisher, catalog #21330) according to the manufacturer’s instructions. The biotinylation reaction incubated overnight at 4°C, after which excess biotin was removed using a 40K MWCO z spin desalting column kit (ThermoFisher, catalog #87766).

### Neutravidin Capture ELISA

30 µL/well of a 0.3 μg/mL solution of neutravidin (ThermoFisher catalog #31000) in ELISA coating buffer was incubated on 384 well microplates at 4°C for 16 hours. The next day, plates were washed 4 times with ELISA wash buffer prior to blocking for 1 hour with 80 µL/well of 1X PBS containing 3% bovine serum albumin and 0.02% Tween 20 (neutravidin blocking buffer). After blocking, plates were washed 4 times with ELISA wash buffer and 30 µL/well of a 2 µg/mL solution of biotinylated gH/gL monomer, 4-mer, 7-mer, 24-mer, or 60-mer was added and allowed to incubate 1 hour. After 4 washes with ELISA wash buffer, a panel of monoclonal antibodies were diluted to 10 µg/mL in neutravidin blocking buffer and three-fold serial dilutions were performed in duplicate followed by a 1-hour incubation at 37°C. 8-16 additional control wells were included that contained immobilized the gH/gL but no monoclonal antibodies (control wells). Following 4 additional washes with ELISA wash buffer, a 1:5000 dilution of goat anti-human IgG (SouthernBiotech, catalog #2010-05) in neutravidin blocking buffer was added to each well and incubated at 37°C for 1 hour followed by 4 washes with ELISA wash buffer. Addition of SureBlue Reserve TMB Microwell Peroxidase substrate, addition of 1N sulfuric acid, and reading of plates was performed as described above.

### Measurement of total plasma IgG

Plasma was serially diluted in ELISA coating buffer in duplicate and incubated on 384-well microplates at 4°C for 16 hours. At least 10 additional control wells were included that contained only coating buffer and no plasma. The next day, plates were washed 4× with ELISA wash buffer prior to blocking for 1 hour with 80 µL/well of ELISA blocking buffer. After blocking, plates were washed 4× with ELISA wash buffer and a 1:4000 dilution of goat anti-mouse IgG (SouthernBiotech, catalog #1030-05) in ELISA blocking buffer was added to each well and incubated at 37°C for 1 hour followed by 4 washes with ELISA wash buffer. Addition of SureBlue Reserve TMB Microwell Peroxidase substrate, addition of 1N sulfuric acid, and reading of plates was performed as described above.

### Bead Depletion Assays

To conjugate biotinylated gH/gL and gH/gL-KO to beads, streptavidin magnetic beads (New England BioLabs, catalog #S1420S) were washed 2× with PBS using a magnetic separator and then co-incubated with biotinylated gp350, gH/gL, or gH/gL-KO on a rotator overnight at 4°C. The supernatant was collected using a magnetic separator and analyzed via spectrophotometry to ensure protein concentration in supernatant had been reduced and saturation of beads was achieved. Beads were washed 2× to remove excess unbound gp350, gH/gL, or gH/gL-KO and stored at 4°C in PBS.

For depletion of plasma antibodies, beads were resuspended with diluted, pooled plasma and incubated 16 hours at 4°C on a rotator. Beads were then separated from plasma using a magnetic separator and the remaining plasma was collected and transferred to a new tube and subsequently tested for binding to gH/gL and for neutralizing activity.

### EBV Challenge in Humanized Mice

12-13 weeks post-human HSPC transfer, 500 µg of total IgG purified from immunized C57BL/6 mice were injected per humanized mouse intraperitoneally (IP). Two days prior to, and one day following transfer, blood was collected to measure the relative levels of total and anti-gH/gL IgG in the plasma.

48 hours after transfer, the mice received a dose of EBV B95.8/F, equivalent to 33,000 infectious units as determined by infection of Raji cells, via retro-orbital injection. Beginning 3 weeks post-challenge (Day 21), peripheral blood samples were collected weekly to determine the presence of EBV DNA in whole blood and to immunophenotype circulating lymphocytes using the antibody panel described in “*Humanized mice.”* Mice were weighed three times a week on non-consecutive days. If mice fell below 80% of their starting weight, or met other criteria for symptoms of pain (i.e., hunching, lack of mobility, etc.), they were euthanized.

Levels of EBV in the blood were monitored on a weekly basis using primers specific for BALF5 as described in “*Quantitative PCR analysis of human cells in huCD34 engrafted mice*.” Blood samples were collected from mice on day prior to challenges and weekly beginning 3 weeks post-challenge (day 21) through to the end of the experiment at 10 weeks post-challenge (day 70), or until the animals reached euthanasia criteria. Spleens were harvested from each mouse at the day 70 endpoint, or earlier if they met euthanasia criteria.

Ten weeks post-challenge, surviving mice were euthanized and spleens were collected and weighed. DNA was extracted from 5×10^6^ total splenocytes, utilizing the DNeasy Blood & Tissue Kit (Qiagen, catalog #69504) and according to the manufacturer’s instructions, for subsequent viral load analysis.

### Quantitative PCR Analysis of Human Cells in HuCD34 Engrafted Mice

A primer-probe mix specific for the EBV BALF5 (Kimura et al., 1999) gene was used to quantify EBV in DNA extracted from blood or spleen in hCD34 engrafted NSG recipient mice at the time points described. Each 25 µL qPCR reaction contained 12.5 µL of 2× QuantiTect Probe PCR Master Mix (QIAGEN), 600nM of each primer, 300nM of FAM-labeled probe, 1.25 µL of a TaqMan 20× VIC-labeled RNase-P primer-probe mix. Reactions were heated to 95°C for 15 minutes to activate DNA polymerase followed by 50 cycles of 95°C for 15 s 60°C for 60 s, on an Applied Biosystems QuantStudio 7 Flex Real Time PCR System. Synthetic DNA fragments containing the BALF5 target gene as well as flanking genomic regions were synthesized as double stranded DNA gBlocks, and were used to generate a standard curve with known gene copy numbers ranging from 10^2^ -10^7^ copies/ml. The copy number in extracted DNA was determined by interpolating from the standard curve. Serial dilutions of reference standard were used to experimentally determine a limit of detection of 6.25 copies, which corresponds to the amount of template that can be detected in >95% of reactions. For graphical purposes, samples with no amplification or those yielding values below the limit of detection were assigned a value of 0.625 copies.

### Biolayer Interferometry

BLI assays were performed on the Octet Red 96 instrument at 30°C with shaking at 1,000 RPM. Anti-Human Fc biosensors (ForteBio, catalog #18-0015) were submerged in wells of black 96-well microplates (Greiner, catalog #655209) containing 250 μL of kinetics buffer (PBS, 0.02%Tween 20, 0.03% azide, 0.1% BSA) for at least 15 minutes prior to any data collection. Biosensors were submerged for 30 seconds in KB to establish baseline response (baseline step 1). Biosensors were submerged in KB containing 10 μg/mL of monoclonal antibodies for 240 seconds (load step). Biosensors were then equilibrated for 60 seconds in kinetics buffer alone (baseline step 2), after which the antibody-bound biosensors were submerged in wells containing a 250 nM solution of gH/gL or gH/gL-KO in KB for 300 seconds (association step) followed by immersion in KB for 300 seconds (dissociation step).

The background signal from each analyte-containing well was measured using empty reference sensors and subtracted from the signal obtained with each corresponding ligand-coupled sensor at every timepoint.

---

**Figure S1.**
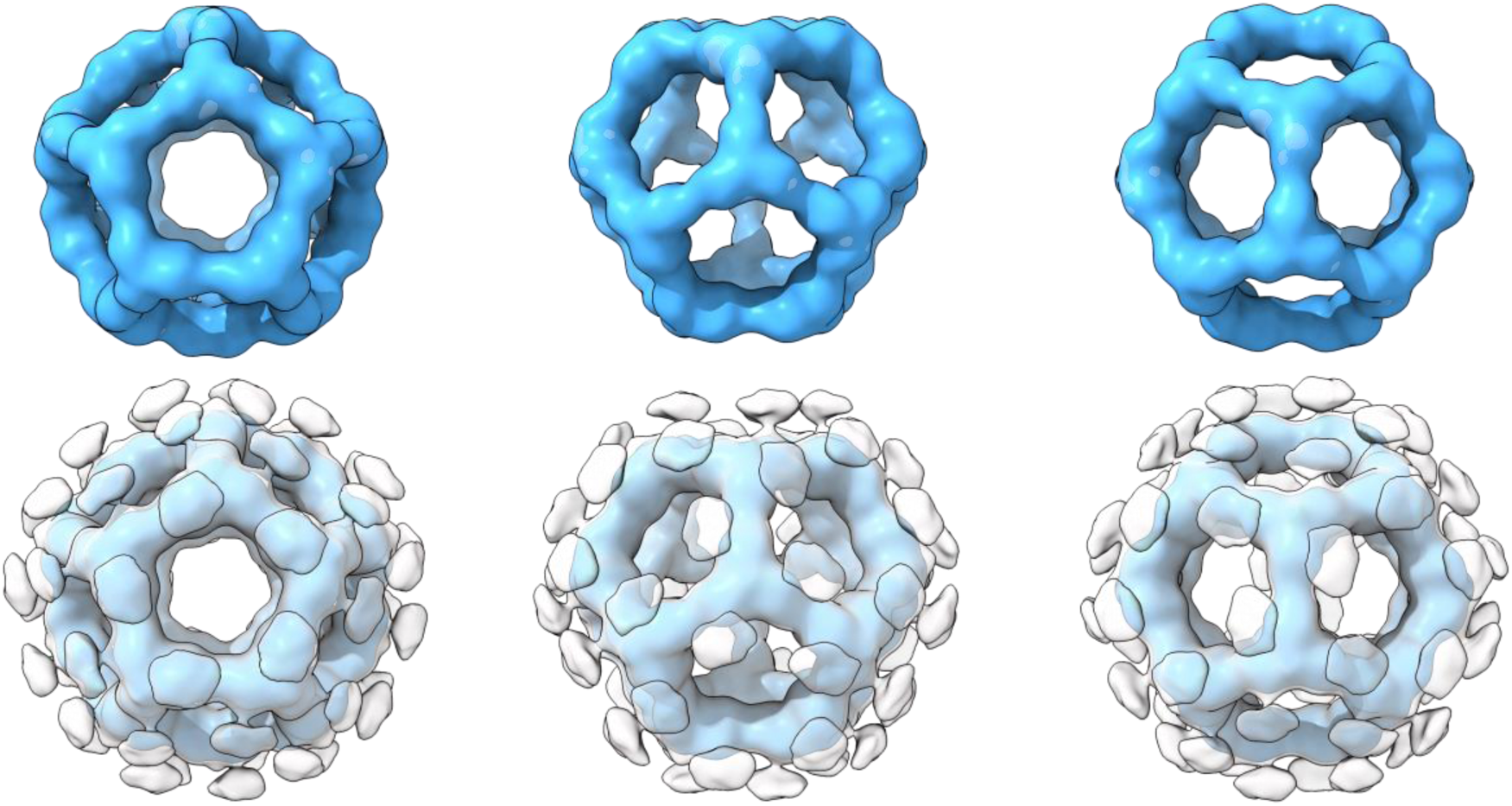
Related to Figure 1. 3D reconstruction of gH/gL-I3 60-mer construct. Negative stain electron microscopy icosahedral 3D reconstruction of the gH/gL-I3 60-mer displayed at different contour levels. (Top) High contour level depicting the I3 nanoparticle base scaffold. (Bottom) Superimposition of two contour levels of the same 3D reconstruction depicting both the I3 base scaffold (blue; high contour) and the presence of displayed flexible gH/gL antigen (grey; low contour). All renderings were generated using ChimeraX.

**Figure S2.**
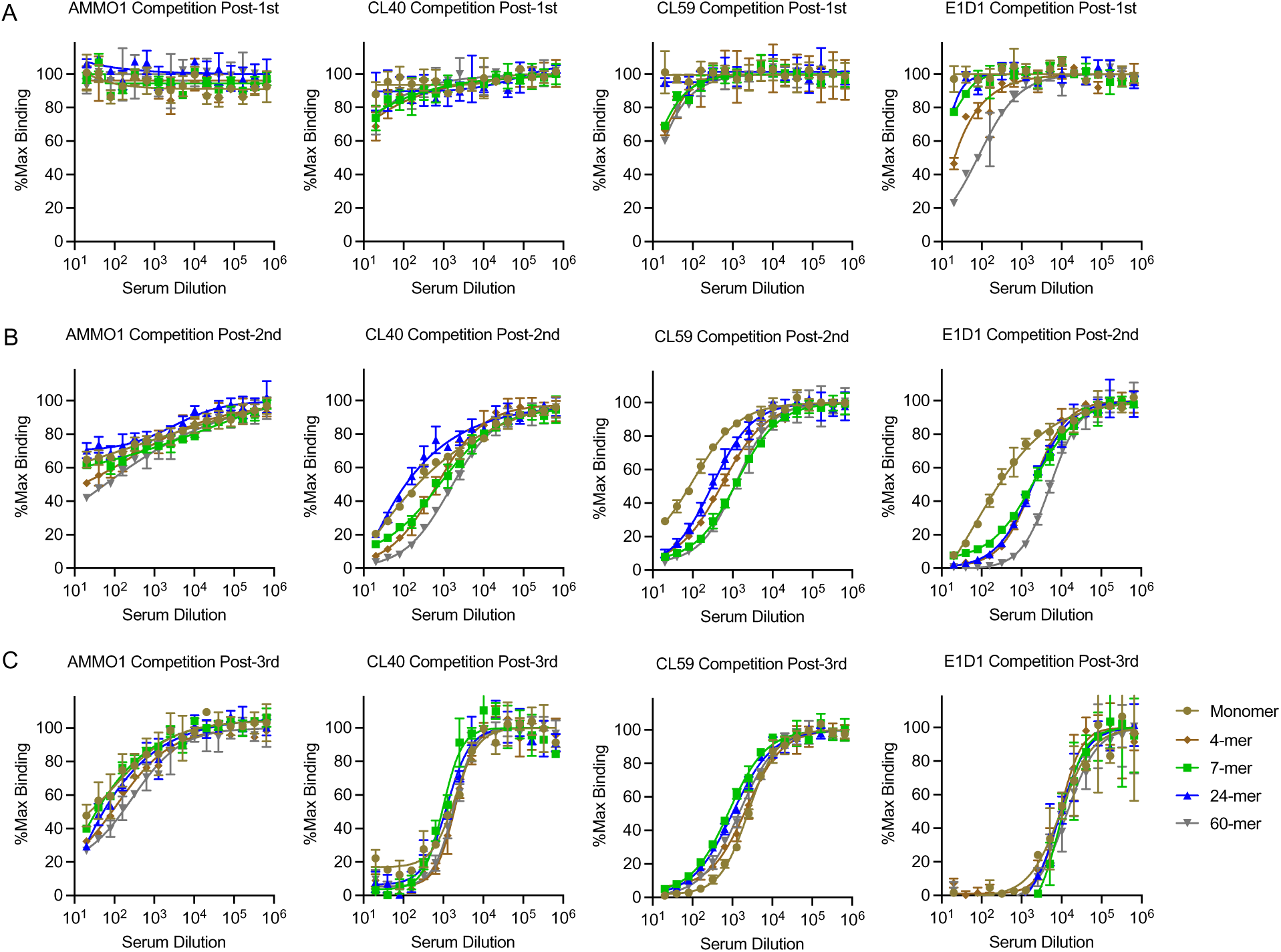
Related to Figure 3. Competitive between immune plasma and monoclonal antibodies. Competitive binding ELISAs were performed using pools of plasma from groups of mice immunized with monomeric gH/gL or multimeric gH/gL nanoparticles, and a panel of anti-gH/gL antibodies. At each time point, pooled sera from each group were titrated on to gH/gL immobilized on an ELISA plate, after which either AMMO1, CL40, CL59, or E1D1 antibodies were added at a concentration previously determined to achieve half maximal binding. Competitions were performed using plasma pools collected at Post-1^st^ (**A**), Post-2^nd^ (**B**), and Post-3^rd^ timepoints (**C**).

**Figure S3.**
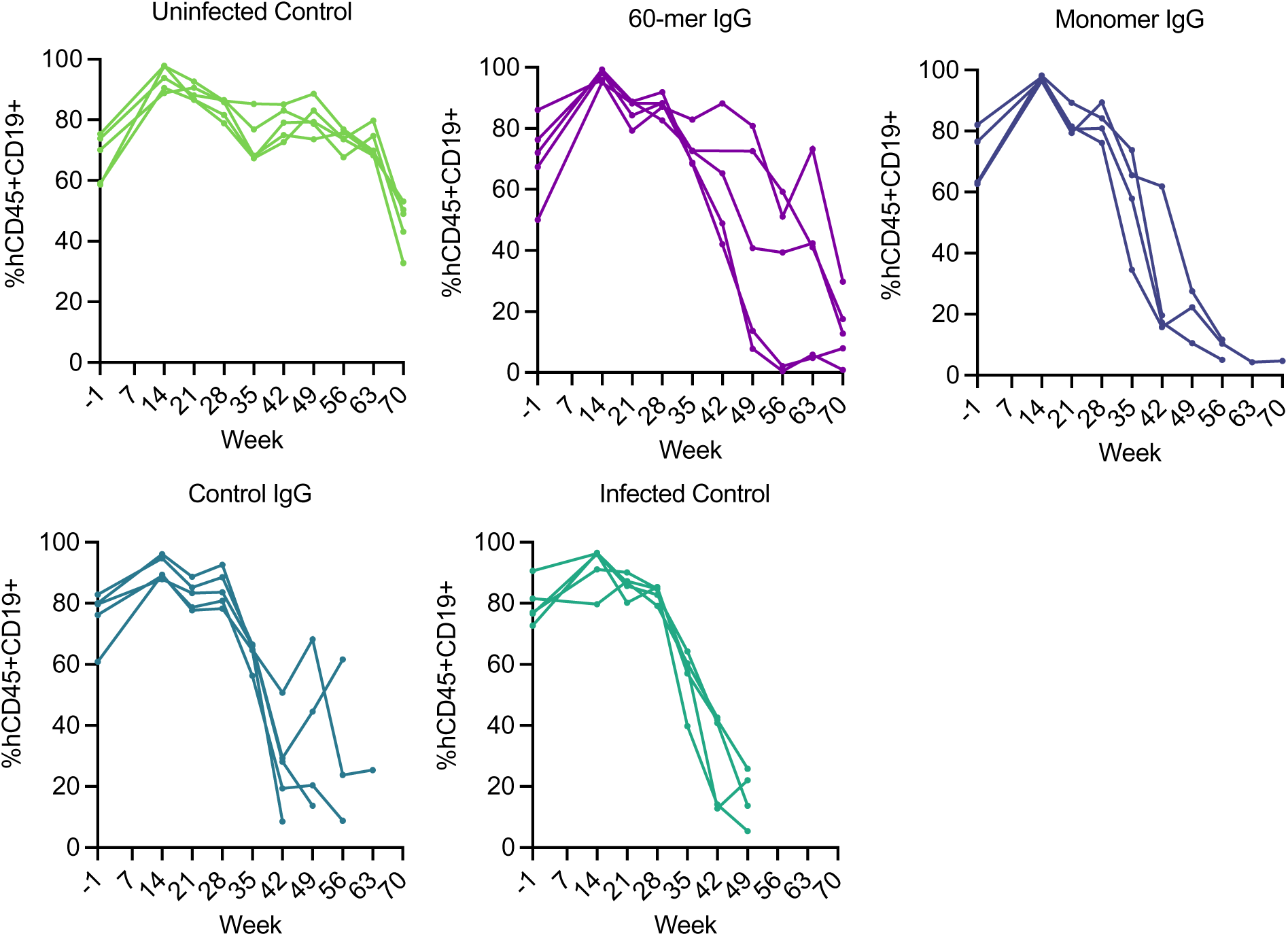
Related to Figure 5. hCD19+ cell frequencies. CD19+ B cell frequencies were measured in the peripheral blood drawn from the mice in Figure 5 at the indicated timepoints via flow cytometry.

**Figure S4.**
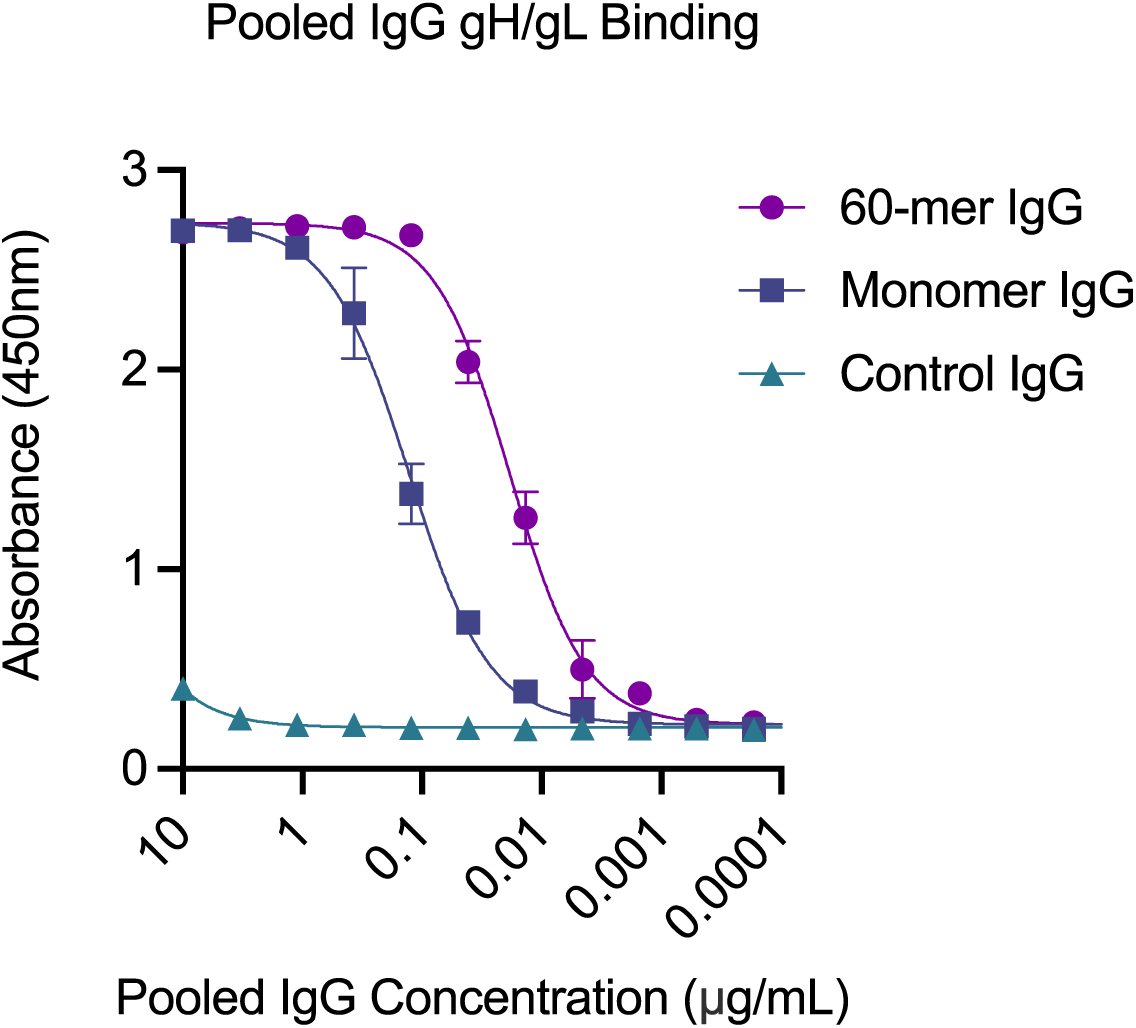
Related to Figure 5. ELISA of pooled purified IgG used in transfer studies. The binding of purified IgG used for adoptive transfer experiments in Figure 5 was measured against gH/gL was measured by ELISA.

**Figure S5.**
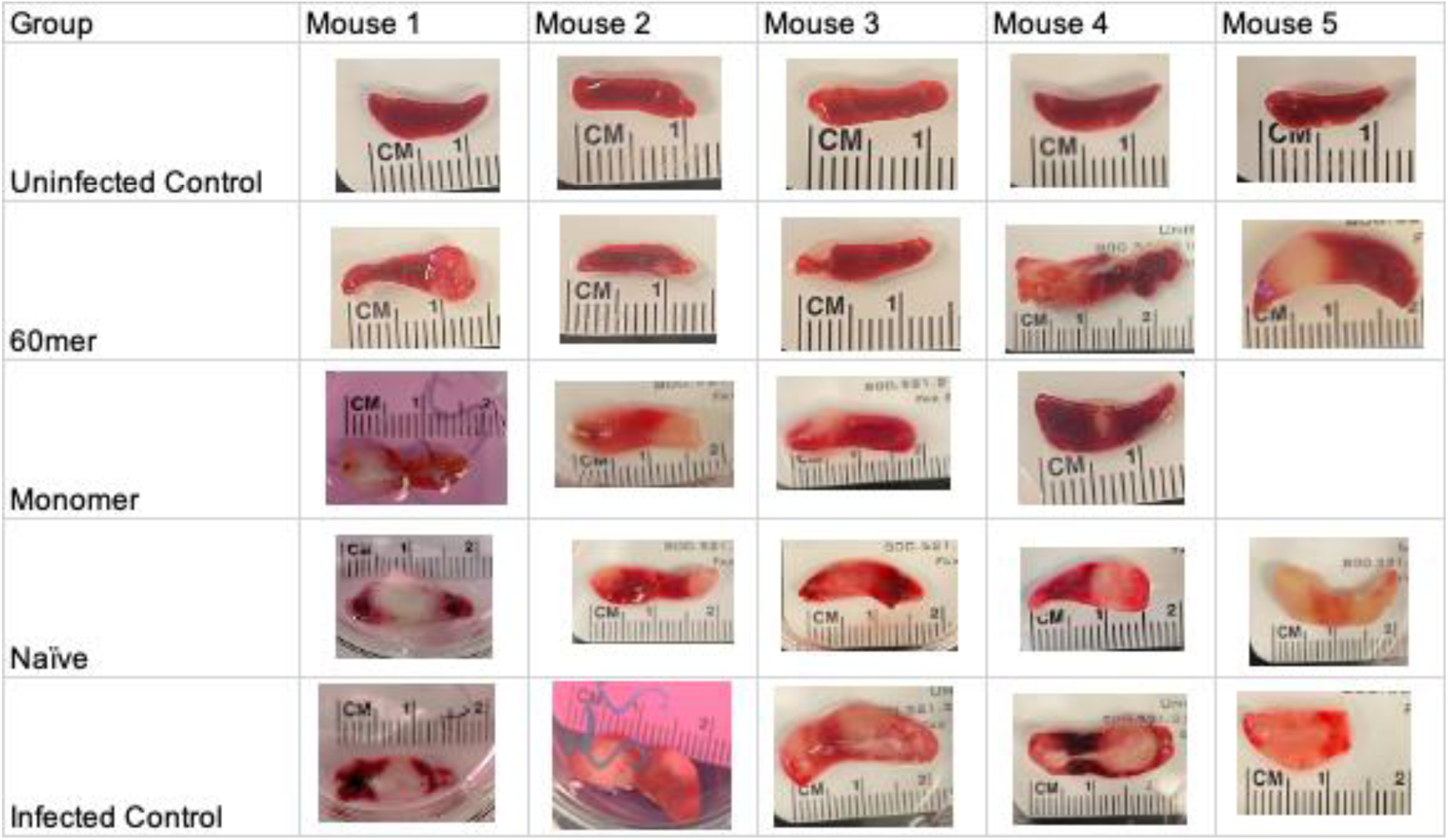
Related to Figure 5. Photographs of spleens from humanized mice challenged with EBV were taken at the time of necropsy.

